# WheatPanRef: a unified linear pan-reference genome for wheat genomics

**DOI:** 10.64898/2026.11.04.764148

**Authors:** Jianxia Niu, Peng Zhao, Wenjie Du, Xiaoyu Zhang, Hang He, Rudi Appels, Jiwen Zhao, Shengwei Ma, Tiantian Chen, Georg Haberer, Manuel Spannagl, Zhanhao Cao, Yuqi Zhang, Lingfeng Miao, Siqi Chen, Baoyue Zhang, Wenxue Wu, Le Wang, Guangwei Li, Jianhui Wu, Zihao Wang, Yongming Chen, Chengzhi Jiao, Xueyong Zhang, Curtis J. Pozniak, Simon G. Krattinger, Frédéric Choulet, Hongqing Ling, Jizeng Jia, Daowen Wang, Wenhao Yan, Zhaorong Hu, Mingming Xin, Huiru Peng, Zhongfu Ni, Surbhi Grewal, Michael Alaux, Rajeev K. Varshney, Weilong Guo, Xing Wang Deng, Qixin Sun

**Affiliations:** State Key Laboratory of High-Efficiency Production of Wheat-Maize Double Cropping, China Agricultural University, Beijing, China; Frontiers Science Center for Molecular Design Breeding, Key Laboratory of Crop Heterosis and Utilization, Beijing Key Laboratory of Crop Genetic Improvement, China Agricultural University, Beijing, China; State Key Laboratory of Wheat Improvement, Peking University Institute of Advanced Agricultural Sciences, Shandong Laboratory of Advanced Agricultural Sciences in Weifang, Weifang, China; Peking-Tsinghua Center for Life Sciences, School of Life Sciences and School of Advanced Agricultural Sciences, Peking University, Beijing, China; Faculty of Science, University of Melbourne, Parkville, Australia; Yazhouwan National Laboratory, Sanya, China; State Key Laboratory for Crop Stress Resistance and High-Efficiency Production, College of Agronomy, Northwest A&F University, Yangling, China; Plant Genome and Systems Biology, Helmholtz Center Munich–German Research Center for Environmental Health, Neuherberg, Germany; Centre for Crop and Food Innovation, Food Futures Institute, Murdoch University, Perth, Australia; State Key Laboratory of Wheat Improvement, College of Agronomy, Shandong Agricultural University, Tai’an, China; College of Plant Science, Jilin University, Changchun, China; State Key Laboratory of High-Efficiency Production of Wheat-Maize Double Cropping, National Wheat Engineering Research Center, College of Agronomy, Henan Agricultural University, Zhengzhou, China; State Key Laboratory of Crop Gene Resources and Breeding, Institute of Crop Sciences, Chinese Academy of Agricultural Sciences, Beijing, China; Crop Development Centre, University of Saskatchewan, Saskatoon, Canada; Plant Science Program, Biological and Environmental Science and Engineering Division, King Abdullah University of Science and Technology, Thuwal, Saudi Arabia; INRAE, GDEC, Université Clermont Auvergne, Clermont-Ferrand, France; Institute of Genetics and Developmental Biology, Chinese Academy of Sciences, Beijing, China; National Key Laboratory of Crop Genetic Improvement, Hubei Hongshan Laboratory, College of Plant Science and Technology, Huazhong Agricultural University, Wuhan, China; Wheat Research Centre, School of Biosciences, University of Nottingham, Loughborough, UK; Université Paris-Saclay, INRAE, BioinfOmics, URGI, Versailles, France

## Abstract

Linear reference genomes provide coordinates for read mapping, variant discovery and multi-omics data integration, yet assembly-specific coordinate systems limit data reuse, particularly in the large, repetitive and polyploid wheat genome. Here, we present WheatPanRef, a scalable linear pan-reference and coordinate framework that integrates wheat sequence diversity within a stable coordinate system. We reconciled independent Chinese Spring assemblies by resolving structural and sequences discrepancies to constructed a 14.50-Gb consensus sequences CS-NP. Using CS-NP as backbone, we incorporated 4.37 Gb of non-redundant non-CS sequences from 37 diverse wheat accessions, generating WheatPanRef (18.87 Gb). WheatPanRef improves read-alignment performance and captures putative wild-relative introgressions and sequences absent or unanchored in conventional references. Bidirectional coordinate-translation system enables accurate projection between WheatPanRef and individual assemblies without remapping, facilitating genomic resource integration and reuse. WheatPanRef provides a common sequence and coordinate foundation for population-scale wheat genomics, with potential utility for genomic resource integration in other complex crops.

## Introduction

Linear reference genomes providing a stable coordinate system for analyzing genomic variation (*1*), read mapping, variant discovery and multi-omics integration (*2*). Long-read sequencing has rapidly expanded the number and completeness of high-quality genome assemblies, accelerating the transition to the pangenome era (*3*). However, each assembly defines its own coordinate system, creating challenges for harmonization, interoperability and reuse of legacy datasets (*1*). Many databases and high-throughput workflows still rely on linear coordinates, making coordinate consistency as important as sequence completeness (*2*). This tension is particularly acute in bread wheat (*Triticum aestivum* L., 2n = 6x = 42), which has a large, highly repetitive hexaploid genome (AABBDD, ∼16 Gb) and is now represented by multiple high-quality assemblies with distinct coordinate systems (*4*).

Reference genome selection has historically reflected community convention and practical utility. Advances in sequencing and assembly have progressively improved reference quality, from chromosome-scale pseudomolecules to near-complete and telomere-to-telomere (T2T) assemblies (*5*). Yet even T2T assemblies may retain sequencing errors, structural or orientation discrepancies. Chinese Spring (CS) has long served as the primary wheat reference, progressing from IWGSC RefSeq v1.0 (*4*) and RefSeq v2.1 (*6*) to the recent near-complete CS-CAU (*7*) and gap-free T2T CS-IAAS assemblies (*8*). These resources represent complementary advances in wheat genomics, while independently generated assemblies can retain distinct strengths and residual discrepancies. Rather than continually replacing references, reconciling assemblies using independent evidence could establish a unified, evidence-weighted consensus backbone formultiple applications.

Reconciling multiple assemblies of the same genotype improves reference accuracy but does not capture diversity among genotypes. A single genotype does not represent the full genomic diversity of the species. Bread wheat has a reticulate evolutionary history and has been extensively enriched by gene flow and alien introgression during evolution and breeding, generating pervasive structural variation, including presence– absence variations (PAVs) and copy-number variations (CNVs) (*9*, *10*). Multi-genome studies have revealed extensive structural divergence among wheat cultivars and associated large-scale structural variation to habitat adaptation and breeding history (*11*, *12*), while recent wheat pangenome analyses have further expanded the catalogue of structural and gene-level diversity (*13*). Sequences absent from or highly diverged relative to a single reference can be missed or incorrectly aligned, resulting in single-reference bias (*1*). Graph-based pangenomes provide a powerful framework for representing alternative alleles, haplotypes and structural configurations (*14*, *15*). However, graph construction, indexing and downstream analysis require specialized data structures and algorithms, whereas many established workflows for population resequencing and multi-omics analysis remain organized around linear coordinates (*16*, *17*). These computational and interoperability constraints are further amplified by the size, repeat content and polyploid complexity of the wheat genome. Consequently, a unified linear pan-reference remains an indispensable tool for wheat genomics, as it must balance comprehensive sequence capture against coordinate stability and computational scalability. Just as the Mercator map projection in cartography distorts continent sizes but preserves local directions for short-distance navigation, we argue that a flat pan-reference optimized for read-mapping provides an accurate coordinate system for genomic intervals up to tens of megabases and will offer the most pragmatic strategy for contemporary wheat genomics and data-driven breeding.

Here, we present WheatPanRef as a consensus-based linear pan-reference for bread wheat. We reconciled independent CS assemblies using complementary sequencing data and validation evidence to construct the near-perfect CS backbone (CS-NP). We then incorporate non-redundant sequence diversity from high-quality hexaploid wheat assemblies into a scalable linear pan-reference designed for high-throughput read mapping and annotation integration. Finally, using WheatPanRef as the common anchor, we established an accurate coordinate-translation scheme linking the pan-reference to individual assemblies. WheatPanRef provides an interoperable framework for integrating genomic diversity and a foundation for community-scale plant genomic research.

## Results

### Multiple-assembly reconciliation yields CS-NP

We resolved sequence discrepancies between two independent high-quality CS assemblies (CS-CAU (*7*) and CS-IAAS (*8*)) by developing an evidence-guided multi-assembly reconciliation framework (**Fig. 1A**). Whole-genome alignment identified 1,786 inter-assembly discrepancies, comprising 1,771 InDels (106.25 Mb), 4translocations (8.24 Mb) and 11 inversions (30.05 Mb) **(table S1)**, which may reflect technical assembly errors or biological variations. Independent short-read (*4*, *6*)), PacBio long-read, and Hi-C datasets of CS were used to resolve these conflicts.

**Fig. 1.**
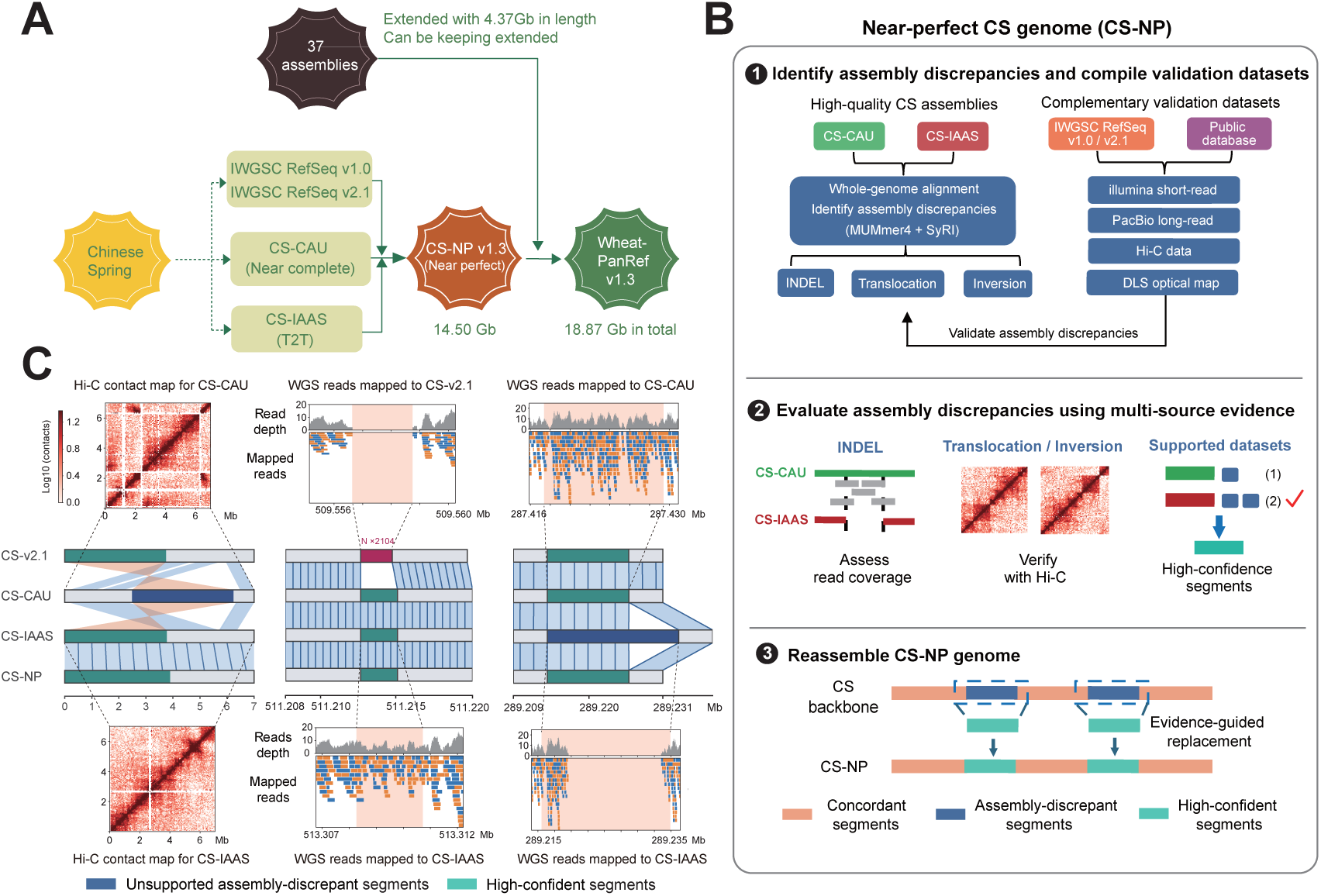
Construction of the CS-NP. **(A)** Overview of CS-NP and WheatPanRef construction. Multiple Chinese Spring (CS) assemblies were integrated to generate the near-perfect Chinese Spring genome (CS-NP). Non-CS sequences identified from 37 additional hexaploid wheat assemblies were subsequently incorporated to generate WheatPanRef. **(B)** Workflow for constructing the near-perfect Chinese Spring genome. Sequence and structural differences among different CS assemblies were identified. The reliability of these regions was evaluated using alignments of Illumina short-read, PacBio long-read, Hi-C and direct label and stain (DLS) optical map datasets. The sequence supported by the greatest number of independent datasets was considered reliable. Poorly supported sequences were replaced with their evidence-supported sequences to generate the final CS-NP assembly. **(C)** Representative examples of evidence-guided reconciliation at assembly-discrepant regions. Syntenic relationships among IWGSC RefSeq v2.1, CS-CAU, CS-IAAS and CS-NP are shown for three regions on chromosome 1B, 6A and 7B. Grey histograms show read depth. Base-collinearity tracks show sequence correspondence among the four assemblies. Pink shading indicates the evaluated intervals. Within the 0–7 Mb region of chromosome 1B, the Hi-C contacts did not support the inversion present in CS-CAU but supported the corresponding sequence arrangement in CS-IAAS, CS-NP therefore adopted the evidence-supported CS-IAAS sequence. On chromosome 6A, a sequence supported by continuous read coverage was incorporated into CS-NP, filling a gap comprising 2,104 Ns in IWGSC RefSeq v2.1. At 289.21-289.23 Mb on chromosome 7B, continuous reads coverage supported the sequence from CS-CAU, which was therefore incorporated into CS-NP.

Discrepancies were evaluated at different genomic scales using the appropriate data type (**Fig. 1B**). Large inversions and translocations were assessed using three independent Hi-C datasets, with contact map across discrepant segments to determine the supported genomic arrangement (**Fig. 1C**). InDels were assessed by comparing short-read coverage across the discrepant segments. Segments with ≥ 80% coverage support were considered reliable. PacBio SMRT long-read provided additional evidence where short-read were inconclusive. For each discrepancy, the sequence supported by more independent datasets was incorporated into CS-NP. Concordant regions were retained unchanged, yielding the initial CS-NP assembly. Further polishing corrected the single-nucleotide errors supported by short-read datasets and resolved medium-scale discrepancies using DLS optical mapping data from IWGSC Refseq v2.1 (*6*). Finally, evidence-guided reconciliation and polishing steps yielded the 14.50-Gb CS-NP assembly **(fig. S1)**. By integrating the sequences best supported by independent datasets, CS-NP provides a unified reference with improved sequence continuity, nucleotide accuracy and structural consistency.

### Iterative framework builds scalable WheatPanRef

To expand the genome sequence representation, we included 37 published chromosome-scale hexaploid wheat assemblies available at the time of dataset compilation **(table S2)**. These assemblies spanned contig N50 values of ∼49 kb to 41 Mb and had BUSCO completeness above 97%. Genome-wide SNP projection showed that these donor assemblies spanned the major diversity space of bread wheat **(fig. S2A).** An independent germplasm resource-based Identity-By-Descent network confirmed the selected genomes represented all nine genetic groups **(fig. S2B)**, supporting their broad genetic representativeness.

To construct the pan-reference while minimizing redundancy, we developed an iterative map-to-pan strategy (**Fig. 2A**), comparing each donor with the current pan-reference rather than all previously incorporated assemblies. This reduced sequence-discovery comparison complexity from quadratic to linear with genome number **(fig. S3)**. To optimize for short-read mapping, at each iteration, unrepresented regions in the current pan-reference were identified as candidate non-CS blocks via a multiscale coverage-depletion scan at resolutions down to 10 Kb, followed by redundancy removal and synteny-based boundary refinement. Cumulatively, all 37 donor assemblies yielded 5,796 non-redundant blocks (4.37 Gb total) spanning all 21 chromosomes **(table S3)**. Projection onto CS-NP revealed uneven homologous positions, often overlapping in distal regions, consistent with multiple haplotypes corresponding to single CS intervals **(fig. S4-6)**. Incorporation of the non-CS pangenome with CS-NP generated the 18.87-Gb WheatPanRef (v1.3).

**Fig. 2.**
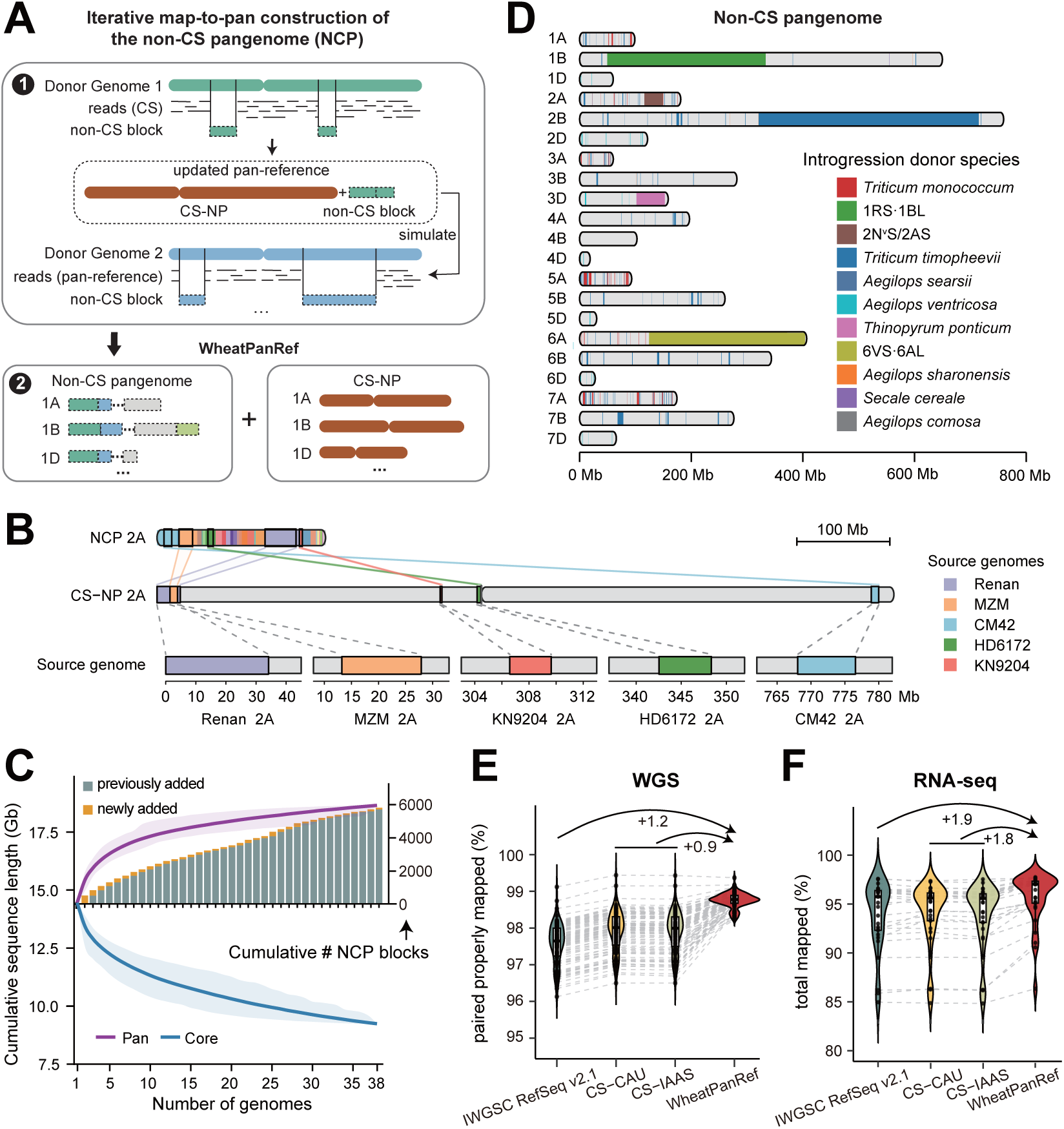
WheatPanRef expands genomic representation and improves read alignment. **(A)** Iterative map-to-pan construction of the non-CS pangenome. Chinese Spring resequencing reads were initially mapped to the first donor assembly, followed by simulated resequencing reads from the expanding pan-reference mapped to successive donor assemblies. Unrepresented non-CS blocks were retained and concatenated into chromosome-specific non-CS pseudochromosomes. **(B)** Positional anchoring and source traceability of non-CS segments on chromosome 2A. Solid lines connect NCP-2A segment to homologous intervals on CS-NP chromosome 2A, and dashed lines link the corresponding CS-NP intervals to their original donor coordinates. Colors indicate donor genomes. **(C)** Pan- and core-genome accumulation across 38 wheat genomes. Curves show means and shaded areas indicate 95% intervals from 1,000 genome-order permutations, with CS-NP fixed first. Stacked bars in the inset show the cumulative previously and newly added non-CS blocks. **(D)** Chromosomal distribution of non-CS sequences assigned to major introgression and putative wild-relative donors. Colored indicate donor classes. **(E, F)** Properly paired read rates for whole-genome resequencing data from 100 wheat accessions (E) and total mapping rates for 29 RNA-seq datasets (F). Grey dashed lines connect measurements from the same sample, arrows indicate mean differences in percentage points.

Importantly, linearization of the non-CS sequences preserved their original genomic context. Local synteny serves as homologous anchors linking retained non-CS segments to the CS-NP genome, while the original donor coordinates were preserved as metadata. Non-CS pseudochromosome 2A (NCP-2A) illustrate three linked coordinate layers: the linearized non-CS pseudochromosome, the corresponding CS-NP interval and the donor assembly (**Fig. 2B**). The Renan segment 2A:0-34.2 Mb contributed to the NCP-2A:116.2-149.5 Mb in the non-CS pangenome, showing synteny to 2A:0-25 Mb of CS-NP genome interval. Such bidirectional tracing links WheatPanRef to CS-NP and source genomes, allowing WheatPanRef to deliver a linear representation of population sequence diversity while facilitating easy retrieval of accurate sequence templates, by retaining the positional context required to locate and interpret alternative haplotypes.

The iterative design also makes WheatPanRef incrementally expandable: new assemblies can be compared directly with WheatPanRef, and non-redundant blocks can be appended or used to refine non-CS regions **(fig. S7)**. Each added genome requires one discovery round rather than comparison with all preceding references or rebuilding the pangenome. This process delivers a version-controlled, efficiently scalable framework for the future expansion of WheatPanRef.

### Expanded genomic representation improves read alignment

Sequence accumulation across the 38 genomes showed that WheatPanRef captured the majority of the sequence diversity whereas the unique contribution of subsequent accession declined progressively. The pan-genome reached 90% of its final size after seven genomes and 95% after 16, while average new sequence per genome declined from 620 Mb for the first three additions to 30 Mb for the last three (**Fig. 2C**). In parallel, the estimated core-genome size progressively decreased and reached 9.24 Gb across the full 38-genome panel. Thus, sequence diversity shared among the currently available assemblies is largely represented in WheatPanRef, whereas continued expansion in future is expected to preferentially contribute rare or lineage-specific sequences.

The expanded non-CS pangenome also captured substantial sequences that were introduced into modern wheat through alien introgression that created chromosome-scale structural variation (**Fig. 2D**). Mapping non-CS blocks to translocation classes and wild-relative genomes identified major breeding-associated introgressions, including 1RS·1BL, 6VS·6AL, 2N^v^S introgression, and also a large *Triticum timopheevii*-derived region on chromosome 2B. In total, 352 non-CS blocks spanning 180.9 Mb were mapped to putative wild-relative donors on the basis of sequence similarity, including 231 blocks assigned to *T. timopheevii*. Thus, WheatPanRef thus captures wild-relative-derived diversity beyond that represented by the single-variety CS reference.

To assess read alignment performance, we compared resequencing data from 100 independent wheat accessions (excluded the 37 donor assemblies) aligned to WheatPanRef and CS references. The WheatPanRef increased total mapping rates by ∼0.1% and simultaneously reduced mismatch rate by ∼0.1% relative to previous CS assemblies **(fig. S8A, B)**. Notably, properly paired read rate increased by 0.9%∼1.2% in WheatPanRef, reaching 98.72% (**Fig. 2E**), corresponding to ∼220 Mb of genomic regions with improved alignment consistency. These gains demonstrate WheatPanRef improves read recovery together with alignment accuracy and consistency.

Across 29 RNA-seq datasets, WheatPanRef increased average mapping rates by 1.9 and 1.8 percentage points over IWGSC RefSeq v2.1 and CS-IAAS, respectively (reaching 95.65%, **Fig. 2F**), with mismatch rates decreased by 0.03 and 0.02 percentage points **(fig. S8C)**. For bisulfite-sequencing (six datasets investigated), mapping efficiency rose by 0.63 percentage points over IWGSC RefSeq v2.1, adding 140 million cytosines for methylation analysis **(fig. S8D, E)**. Thus, WheatPanRef currently serves as the optimal reference for short-read alignment, enhancing the mapping of multi-omics data at population level.

### Multi-source integration delivers consensus annotation

Given the gene structure annotation quality lags substantially behind genome assembly quality, we sought to construct a unified gene catalogue for WheatPanRef by leveraging recurrent support across independent genome annotations. This process was based on the premise that random annotation errors rarely coincide at the same locus. We integrated 38 transferred annotation sets from the assembled wheat genomes and three independently generated sets from international expert groups (**Fig. 3A**). Overlapping models were consolidated into common gene loci. Transcript models sharing the same intron chain were grouped first, and then representative models were prioritized based on cross-source support, transcript completeness and structural consistency (**Fig. 3A**). The final annotation comprised 232,140 gene loci, including 182,360 on CS-NP and 49,780 on non-CS pangenome. WheatPanRef gene identifiers also cross-references legacy *TraesCS* identifiers and identifiers from other wheat assemblies, enabling bidirectional querying of corresponding gene IDs through the WheatPanRef web portal (https://wheat.cau.edu.cn/WheatPanRef/).

**Fig. 3.**
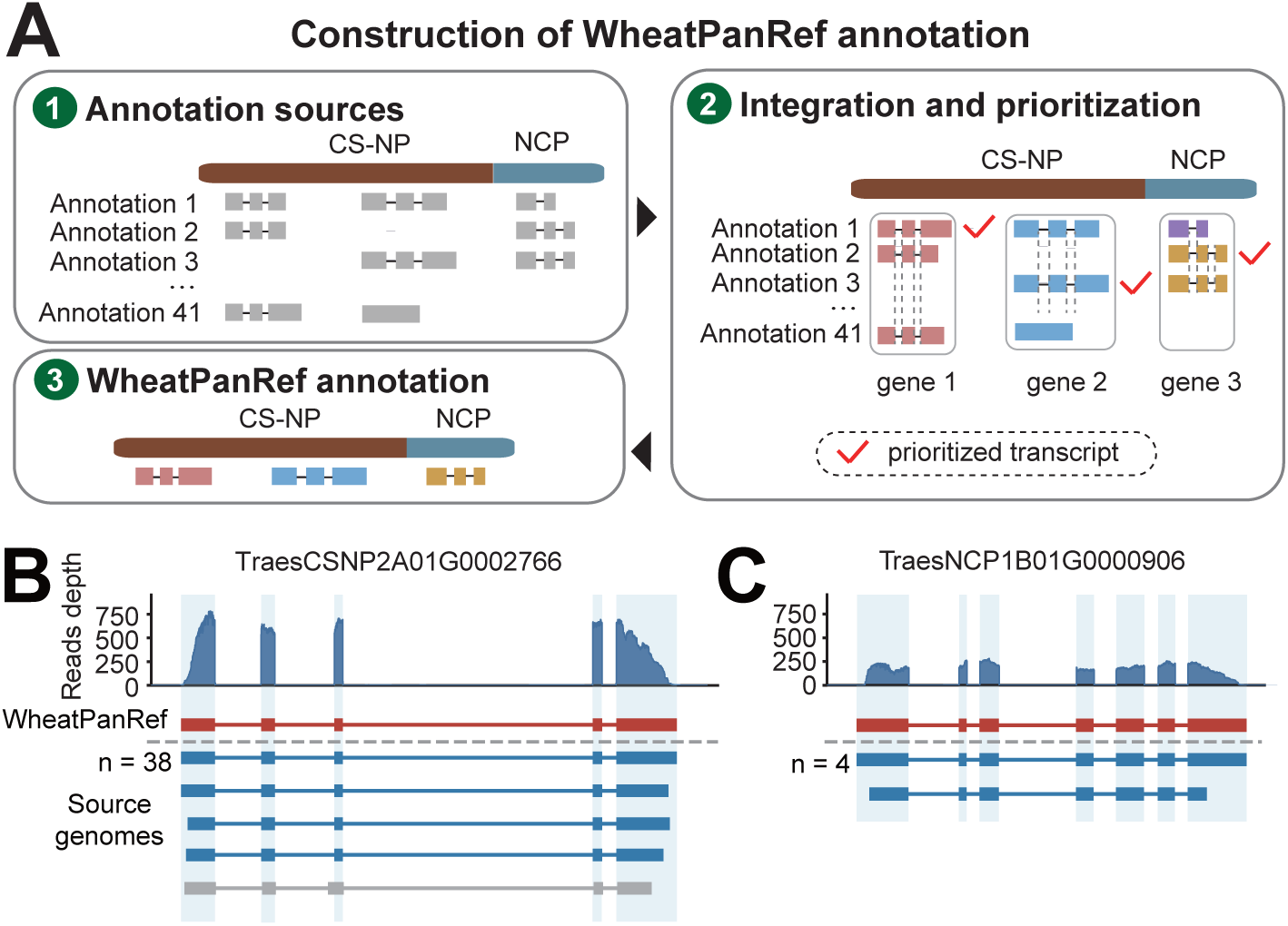
Integrated gene annotation of WheatPanRef. **(A)** Integration of 41 annotation sets. Overlapping gene models were consolidated into loci, and transcript structures were grouped by intron chains for model prioritization. **(B, C)** Representative transcript-model integration at TraesCSNP7A01G0002766 (B) and TraesNCP1B01G0000906 (C). Red models indicate selected WheatPanRef transcripts. Blue and grey models indicate alternative annotation models. *n* denotes the number of supporting annotation sets.

To evaluate the robustness of the integrated annotation, we quantified cross-source support for each transcript structure. Of 238,377 WheatPanRef transcripts, 223,612 (93.8%) were supported by at least two genome annotations, including 81,757 supported by more than 20 sets **(fig. S9A)**. The latter comprised 80,773 CS-NP transcripts and 984 non-CS-genome transcripts, indicating extensive independent support for a substantial fraction of the integrated annotation. Cross-genome recurrence and RNA-seq evidence jointly supported transcript-model prioritization (**Fig. 3B, C**). In both CS-NP and non-CS segments, selected WheatPanRef models showed recurrent transcript structures and exon-resolved RNA-seq support (**Fig. 3B, C**). These results show that the integration framework preferentially retains reproducible, transcriptionally supported gene models. The final WheatPanRef annotation showed high completeness by BUSCO, with 99.7% complete orthologues, comparable to or exceeding the evaluated wheat reference annotations **(Fig. S9B)**. Together, these data indicate that the integration procedure expanded gene representation without compromising annotation quality.

### WheatPanRef enables cross-genome coordinate integration

Assembly-specific coordinate systems scattered genomic features and limited direct comparison and reuse data-mapping results across references, therefore we established a cross-genome coordinate-conversion framework by bidirectionally linking WheatPanRef with other wheat assembly via chain files (**Fig. 4A**). We compared direct coordinate conversion from ten source assemblies to WheatPanRef, Fielder and IWGSC RefSeq v2.1, and found that WheatPanRef achieved improved coordinate conversion rate (mean=97.58%), exceeding the other single-variety references by 5.9 and 7.3 percentage points, respectively (**Fig. 4B**). Meanwhile, the coordinate accuracy (98.52%) was also marginally elevated (**Fig. 4C**). Together, these improvements indicate that the broadened sequence representation of WheatPanRef increases locus recoverability and coordinate mapping capacity.

**Fig. 4.**
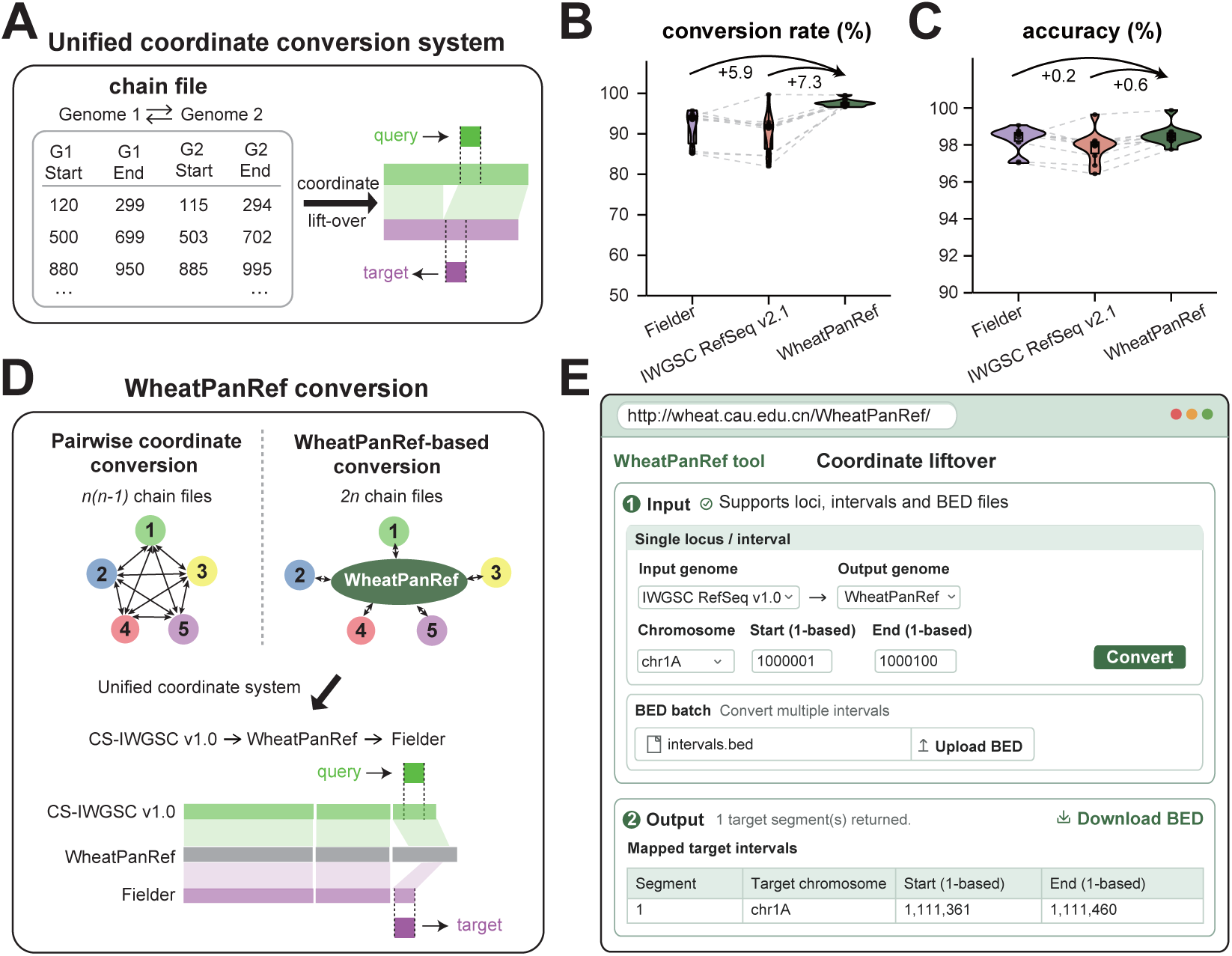
Cross-genome coordinate integration with WheatPanRef. **(A)** Directional chain files linking homologous genomic intervals between assemblies for coordinate conversion. **(B, C)** Direct coordinate conversion from ten wheat assemblies to Fielder, IWGSC RefSeq v2.1 and WheatPanRef, assessed by conversion rate (B) and accuracy (C). Ten sets of 1,000 loci were sampled independently from each of four chromosomes per assembly. Black dots show assembly-level means. Grey dashed lines connect the same source assemblies. Arrows indicate mean differences in percentage points. **(D)** Pairwise versus WheatPanRef-mediated coordinate conversion strategies. Linking *n* assemblies bidirectionally through WheatPanRef requires *2n* directional mappings, compared with *n(n − 1)* pairwise mappings. The lower schematic shows cross-genome conversion through WheatPanRef. **(E)** Web interface for converting genomic intervals and BED files between assemblies linked through WheatPanRef.

Rather than requiring exhaustive pairwise mappings a shared coordinate system through WheatPanRef enables scalable cross-genome conversion between individual assemblies (**Fig. 4D**), dramatically reducing the number of required mappings coordinate system complexity from quadratic to linear. Moreover, WheatPanRef also provides an intermediate reference for inter-genome coordinate conversion, preserving recoverability and positional accuracy, while increasing the mean conversion rate by 3.2 percentage points over standard single references and maintaining comparable accuracy **(fig. S10)**. These results demonstrate that WheatPanRef serves as a central hub in the coordinate layer, enabling accurate conversion among multiple assembly-specific coordinate systems with high cross-genome transferability.

To make the coordinate framework directly accessible, we developed a web-based interface for coordinate conversion between WheatPanRef-linked assemblies (**Fig. 4E**). Downloadable chain files are also provided to enable offline coordinate transfer **(fig. S11)**. Together, these resources facilitate the reuse and integration of genomic data across wheat reference systems.

### WheatPanRef captures genomic diversity beyond single genomes

We next asked what types of gene content and functional diversity were preferentially represented in sequence captured beyond the CS-NP backbone. Notably, NLR loci were distributed across all 21 chromosome-specific non-CS components, including regions corresponding to major breeding-associated introgressions such as 1RS·1BL, 2N^v^S introgression (from *Aegilops ventricosa*), 6VS·6AL (from *Dasypyrum villosum*) and segments derived from *Triticum timopheevii* and *Thinopyrum ponticum* (**Fig. 5A**). Several well-characterized resistance loci, including *Pm8*(*18*), *Yr9*(*19*), *Lr13*(*20*), *Pm6/Pm52*(*21*), *Pm41*(*22*) and *Pm21*(*23*), were represented within these non-CS components. Across WheatPanRef, we identified 2,306 non-redundant NLR clusters, of which 1,410 (61.1%) were located on the non-CS genome and 896 (38.9%) on CS-NP (**Fig. 5A**). After accounting for sequence length, NLR-cluster density was approximately fivefold higher in the non-CS pangenome than on the CS-NP backbone, demonstrating that non-CS sequence substantially expands the representation of clustered immune-receptor diversity. Consistent with this pattern, Gene Ontology analysis revealed a modest enrichment of defense-response genes in the non-CS pangenome (1.20-fold enrichment, **fig. S12**). Although the magnitude of enrichment was modest, these results independently support an increased representation of defense-associated genes in the non-CS pangenome.

**Fig. 5.**
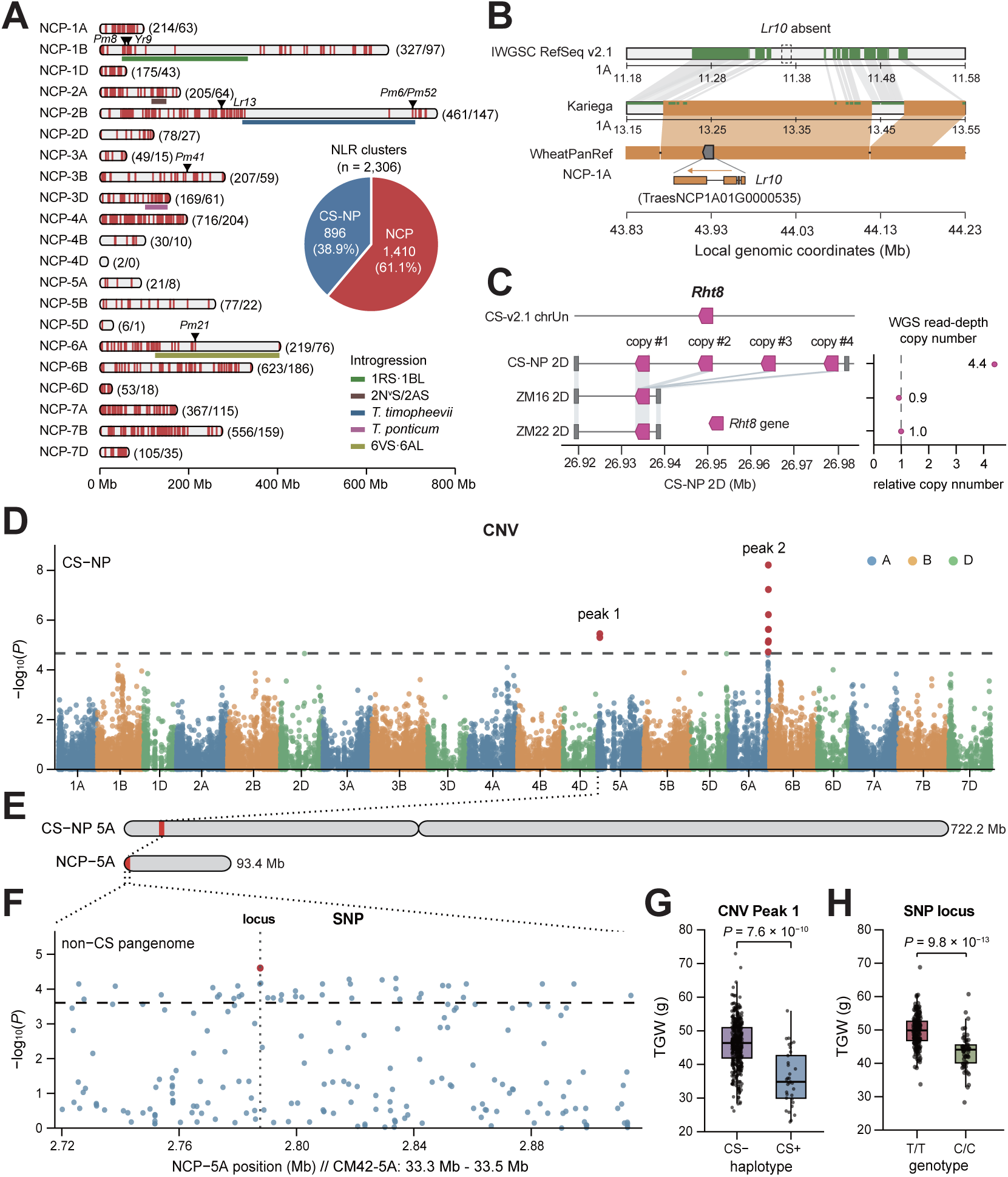
Functional diversity captured by WheatPanRef. **(A)** Distribution of NLR cluster and major introgressions across non-CS chromosomes. Red marks indicate NLR cluster. Colored intervals denote introgression classes. Numbers in parentheses indicate NLR locus and cluster counts. **(B)** Recovery of *Lr10*, absent from IWGSC RefSeq v2.1, in a Kariega-derived chromosome 1A segment incorporated into NCP-chr1A. **(C)** Copy-number variation at *Rht8* locus. Pink arrows indicate *Rht8* copies, and grey lines indicate sequence homology. **(D)** CNV association analysis for thousand-grain weight (TGW) in 485 wheat accessions. Red points indicate associations exceeding the suggestive significance thresholds (dashed line). **(E)** Correspondence between the chromosome 5A deletion-associated interval in CS-NP and the CM42-derived alternative sequence in NCP-5A. **(F)** Local SNP association analysis within the CM42-derived sequence. The red point indicates the lead SNP, and the dashed line indicates the region-specific Bonferroni significance threshold. **(G)** Thousand-grain weight (TGW) distributions of accessions carrying the deletion-associated NCP haplotype (DEL/NCP+) or the CS-NP reference haplotype (REF/NCP−) at the chromosome 5A CNV-associated locus. **(H)** TGW distributions of the two genotypes (T/T and C/C) at the lead SNP within the CM42-derived alternative sequence. P-values in (G) and (H) were calculated using two-sided Wilcoxon rank-sum tests.

We next asked whether the expanded sequence space of WheatPanRef could provide genomic context for functionally important variation that is absent from or incompletely represented in conventional CS references. The leaf-rust resistance gene *Lr10*, absent from IWGSC RefSeq v2.1, was recovered within a Kariega-derived 1A segment incorporated into NCP-1A, where it is represented by TraesNCP1A01G0000535 (**Fig. 5B**). By contrast, the dwarfing locus *Rht8* (TraesCSU03G0022100), which resides on chrUn in IWGSC RefSeq v2.1, was anchored to CS-NP chromosome 2D and resolved as a four-copy tandem array at ∼28.01-28.07 Mb (**fig. S13**). Comparative analysis identified single-copy *Rht8* configurations in ZM16 and ZM22, consistently supported by genome assemblies and resequencing read-depth analysis (**Fig. 5C**). Together, these examples show that WheatPanRef recovers missing or incompletely represented functional variation and places it within sequence-resolved genomic contexts for downstream trait mapping.

To determine whether WheatPanRef could resolve trait-associated variation beyond the CS-NP reference, we performed CNV association analysis for thousand-grain weight (TGW) in 485 wheat accessions. A significant deletion-associated peak on CS-NP chromosome 5A was detected (*P* = 3.54 × 10⁻⁶), corresponding to a CM42-derived alternative sequence fragment in WheatPanRef, spanning 990.6 Kb and harboring 14 annotated genes (**Fig. 5D, E and fig. S14**). A total of 447 accessions carrying the fragment-deletion showed significantly decreased TGW compared to wild type (*P* = 7.6 × 10⁻^10^; **Fig. 5G**). We further examined the GWAS results by tracing to the CM42-derived sequences and identified 39 significantly associated SNP loci (**Fig. 5F**). The most significant SNPwere further partitioned into two genotypes, with the T/T genotype showing significant higher TGW than C/C (*P* = 9.8 × 10⁻¹³; **Fig. 5H**). Together, these results demonstrate that WheatPanRef supplies a reference template for SV-based GWAS and enables fine-scale haplotype discovery for trait-associated alternative alleles.

## Discussion

Despite advances from chromosome-scale assembly to T2T gap-free CS assembly (*6–8*), sequence and structural discrepancies persist among independently assemblies. By integrating multiple CS assemblies using complementary evidence, we constructed a unified and refined CS genome (CS-NP) that resolves assembly errors and distinguishes true polymorphisms from assembly artifacts. This CS-NP genome minimizes false-positive variant detection associated with single-assembly biases, consolidating the strengths of multi-assemblies into a community standard reference. The proposed evidence-guided reconciliation strategy extends genome improvement beyond completeness to cross-assembly consistency, offering a framework for other plant varieties.

WheatPanRef expanded the CS backbone with non-redundant fragments from diverse assemblies, strengthening the ability to capture population-level diversity beyond any single reference genome, with improved short-read mapping rates and reduced mismatch rates. Unexpectedly, the non-CS genome contain numerous wild-relative-derived regions and are ∼5-folder enriched of *NLR* genes relative the CS genome, underscoring how introgression and an extensive NLR reservoir have shaped wheat adaptation, and responses to evolving pathogen pressures in wheat breeding (*11*, *12*). Morevoer, recovering *Lr10* and resolving new TGW-QTL locus illustrate how WheatPanRef retrieves variation ‘missing’ in a CS-only framework and permits further haplotype interrogation of deletion-associated traits. The iterative map-to-pan strategy for expanding and updating the pan-reference enables continued addition of new assemblies with minimized comparisons efforts, and will serve as a scalable framework for community-wide pan-genome construction amid the arrival of hundreds to thousands of incoming assemblies.

As a common coordinate bridge, WheatPanRef enables reuse of legacy genomic and multi-omics datasets through bidirectional coordinate mapping, reducing the efforts to benchmark historical resources against each new reference. Its broad sequence representation enables the pan-reference to act as the intermediate system for cross-genome conversion. Crucially, continued compatibility with established linear-reference workflows keeps this a practical operational strategy for current pipelines, and the same coordinate system is well positioned to interface with emerging graph pangenomes through coordinate translation. Collectively, WheatPanRef delivers a unified, expandable foundation for wheat genomics, and is well positioned to streamline multi-omics integration at scale in the pan-genomic era. Beyond wheat, the logic of reconciling multiple assemblies and anchoring dynamic sequence diversity to a stable coordinate axis offers a transferable blueprint to operationalize consensus coordinate system for the broader crop genomics community.

## Materials and methods

### Identification of discrepancies between CS assemblies

The near-complete CS assembly (CS-CAU) (*7*) was aligned chromosome by chromosome to the telomere-to-telomere CS assembly (CS-IAAS) (*8*) using MUMmer (v4.0) (*24*) with parameters: ‘--maxmatch -g 300 -c 200 -l 500’. Alignments were filtered using delta-filter with parameters: ‘-m -r -q -i 90 -l 500’, and aligned results were processed using ‘show-coords -THrd’ to obtain the coordinate files. Assembly discrepancies were identified using SyRI (v1.6) (*25*) with default parameters and classified as insertions, deletions, inversions and translocations.

### Evaluation of assembly discrepancies using multi-source evidence

The Illumina whole-genome sequencing (WGS) reads from three independent CS libraries were aligned to CS-IAAS and CS-CAU using BWA-MEM (v0.7.15) (*26*) **(table S4)**. PacBio SMRT long-read from the CS were aligned to CS-IAAS and CS-CAU using minimap2 (v2.3) (*27*) with the PacBio preset (-x map-pb -t 12 -L). Only alignments with a mapping quality (MAPQ) ≥ 20 were retained. For each insertion or deletion, read coverage and mean read depth were calculated across the interval and its 50-bp flanking regions. Intervals with read coverage of at least 80% and a mean read depth of at least 2× were considered reliably supported. For each discrepancy, sequence segments supported by more independent datasets (at least two datasets) were classified as high-confidence sequence and selected for CS-NP construction. When neither sequence was supported by at least two short-read datasets, an independent PacBio SMRT long-read dataset was used as additional evidence. When both sequence segments were supported by the same number of datasets, read coverage and read depth across the corresponding intervals were compared to select the sequence with stronger read support. Assembly-specific intervals, which were represented in only one assembly, were considered reliable when supported by Illumina dataset or the PacBio dataset.

Hi-C reads from three independent CS libraries were aligned to CS-IAAS and CS-CAU using chromap (v0.3.2) (*28*) with the Hi-C preset. PCR duplicates were removed, and read pairs with a minimum mapping quality of 1 were retained (--preset hic --remove-pcr-duplicates -q 1). Contact maps were subsequently generated using juicer_tools (v1.9.9) (*29*). The contact maps were visualized in Juicebox (v1.11.08) (*30*) and used to evaluate contact continuity across inversion and translocation regions. For each inversion or translocation, the corresponding Hi-C contact maps generated for CS-IAAS and CS-CAU were compared across the three independent datasets. The assembly configuration displaying more continuous near-diagonal contacts and fewer breakpoint-associated off-diagonal contacts was considered better supported. A structural configuration was selected when it was consistently supported by at least two of the three Hi-C datasets.

### Construction of the CS-NP assembly

Sequence regions concordant between the CS-CAU and CS-IAAS assemblies were retained unchanged. At each assembly discrepancy, the alternative sequence supported by the independent WGS, PacBio or Hi-C evidence described above was selected and incorporated at the corresponding genomic position, thereby generating the initial CS-NP assembly. To correct residual single-nucleotide errors, Illumina WGS reads from three independent CS datasets were aligned to the initial CS-NP assembly using BWA-MEM (v0.7.15) (*26*). SNPs were called using GATK HaplotypeCaller (v3.8.1) (*31*). SNPs with the genotype quality (GQ) ≥ 30 and read depth ≥ 2 were retained. A reference base was replaced when the same alternative allele was supported by at least two of the three independent WGS datasets. Three independent CS Hi-C datasets were then aligned to the corrected CS-NP assembly. The resulting contact maps were manually inspected for potential chromosome-scale misassemblies, including discontinuities in near-diagonal contacts and anomalous off-diagonal signals. Finally, direct label and stain (DLS) optical map (*6*) generated for CS was aligned to the Hi-C-validated CS-NP assembly for further structural refinement. Regions showing discordant optical-map alignments, including unsupported gaps, overlapping alignments and orientation inconsistencies, were manually examined. Regions identified as potential misassemblies were corrected when supported by the DLS optical map and complementary sequencing evidence. The resulting assembly constituted the final CS-NP reference.

### Iterative construction of WheatPanRef

To expand genomic representation while minimizing sequence redundancy, we developed an iterative map-to-pan strategy. A total of 37 chromosome-scale hexaploid wheat assemblies (*11*, *12*, *32–41*) were incorporated sequentially, prioritized by contig N50 **(table S2)**. For the first donor (CM42), Illumina whole-genome sequencing (WGS) reads from Chinese Spring (CS) were preprocessed with fastp (*42*) and aligned to CM42 using BWA-MEM (*26*). Alignments were processed with SAMtools (*43*) and BamTools (*44*), retaining reads with mapping quality ≥1 and non-zero absolute insert sizes <10 kb. To capture non-CS intervals across different genomic scales, we performed a multiscale coverage-depletion scan using BEDTools (*45*) across nine window sizes (1 Mb, 500 kb, 200 kb, 100 kb, 50 kb, 40 kb, 30 kb, 20 kb, and 10 kb) with a 20% step size. We normalized the read density of each window against the genome-wide background read depth (defined as the mode of non-zero read density). Windows with a normalized density ≤0.5 were merged to define candidate intervals. Large candidates were tracked from the 1-Mb to finer resolutions, selecting the preceding coarser resolution when boundary displacement approached the minimum within 50 kb.

Candidate blocks were filtered for sequence redundancy using MUMmer4 (*24*). Sequences with an identity-weighted aligned fraction >30 to the CS-derived sequence or previously retained blocks were removed. We refined the boundaries of candidate blocks through local synteny analysis. Each candidate interval was extended by 2 Mb on both sides within its donor assembly and aligned to the corresponding CS chromosome using MUMmer4 (*24*). We merged collinear intervals and used the complementary non-collinear intervals overlapping the original candidate to redefine its limits. Retained blocks were concatenated into 21 chromosome-specific non-CS pangenomes, separated by 500-bp stretches of Ns, and combined with the CS-NP chromosomes to generate WheatPanRef. From the second iteration onwards, we simulated 150-bp paired-end reads from the cumulative pan-reference pool using GenerateReads_WG_PE and mapped them to each subsequent donor assembly to identify novel specific blocks via the same multiscale coverage-depletion procedure.

### Pan- and core-genome accumulation analysis

To quantify the diversity captured by the expansion, we partitioned WheatPanRef into 1-Mb windows (with non-CS blocks partitioned independently to avoid artificial junctions). We aligned each window to the corresponding chromosomes of the 37 donor assemblies using minimap2 (*27*) (asm5 preset) and considered a window present if ≥30% of its non-N sequence aligned at ≥80% nucleotide identity. Pan- and core-genome accumulation was assessed using 1,000 random permutations of the 37 donors, with CS-NP fixed as the first genome. The pangenome and core genome were defined as the union and intersection of present windows, respectively. The mean values and 95% confidence intervals were calculated based on these random permutations.

### Sequence-based assignment of non-CS blocks to putative wild-relative donors

To trace the origins of non-CS diversity, we compared non-CS blocks against genome assemblies of 11 wheat wild relatives, including *Aegilops comosa* (*46*), *Aegilops longissima* (*47*), *Aegilops searsii* (*47*), *Aegilops sharonensis*(*47*), *Aegilops ventricose* (*48*), *Aegilops tauschii* (*49*), *Dasypyrum villosum* (*50*), *Secale cereale* (Lo7 (*51*) and Weining (*52*)), *Thinopyrum elongatum* (*53*), *Triticum monococcum* (*54*) and *Triticum timopheevii* (*55*) **(table S5)**. Each block was aligned to the corresponding wild-relative chromosomes using MUMmer4 (*24*) (nucmer --mum -c 100 -l 50), followed by delta-filter (-o 90 −1). To minimize interference from dispersed repetitive alignments, we restricted scoring to the five best-supported 3-Mb windows (scanned at 100-kb steps). Sequence similarity was calculated as the cumulative non-redundant aligned length multiplied by mean nucleotide identity (%), divided by block length. A wild-relative assembly was assigned as the putative donor if the similarity score exceeded 70.

### Integration and quality control of WheatPanRef gene models

To overcome annotation fragmentation and random errors, we integrated 41 complementary annotation sets, based on the premise that independent annotation errors rarely coincide at the same locus. Existing annotations were projected using Liftoff (*56*) with minimap2 (*27*), requiring sequence identity ≥ 95% and alignment coverage ≥ 60%. In addition, we incorporated three independently generated annotation sets from international expert groups. The first set (by Manuel group) used a projection-based approach, transferring high-quality gene predictions from established wheat references (IWGSC RefSeq v1.1 and gene predictions from the 10+ Wheat Genome Project) using a non-redundant gene projection pipeline (*57*). The second set (by Shengwei group) was produced using the NCBI Eukaryotic Genome Annotation Pipeline (EGAPx v0.4.1) after screening the WheatPanRef assembly with the NCBI Foreign Contamination Screen (*58*). This annotation integrated short- and long-read RNA-seq evidence together with protein evidence from related Poales species, and its completeness was evaluated using BUSCO(*59*). The third set (by Yongming group) assembled transcripts from RNA-seq reads unmapped to IWGSC RefSeq v2.1 using Trinity, followed by redundancy removal (CD-HIT, 90% identity) and contaminant filtering (DeconSeq/BLASTN). These transcripts were aligned to WheatPanRef using GMAP (*60*) (≥95% identity), and integrated with TransDecoder-predicted coding sequences.

We consolidated the 41 annotation sets using a hierarchical procedure to define non-redundant gene loci and representative transcript models. Initially, gene models located on the same chromosome and strand with overlapping genomic intervals were grouped into provisional loci. Within each locus, we clustered transcripts with identical exon-intron structures. Specifically, multi-exon transcripts were grouped if they shared the same number of exons and the boundaries of all corresponding introns differed by ≤10 bp. Single-exon transcripts were grouped if they exhibited at least 50% reciprocal overlap. For each transcript cluster, we calculated a support score based on the number of independent annotation sets that contributed to it. The intron-chain structure with the highest support score was selected as the primary isoform. When multiple transcripts shared this primary structure, we retained the one with the greatest total exon length, using source priority, sequence identity, alignment coverage, and ORF integrity to resolve any remaining ties. Following selection, we applied structural and protein-level quality control by correcting CDS phases with AGAT, extracting sequences via gffread (*61*), and excluding transcripts containing internal stop codons. Ultimately, for each gene locus, the transcript models with the highest support score were retained as the final, high-confidence gene annotation set for WheatPanRef.

### Construction of cross-genome liftover chains

To enable bidirectional coordinate translation, we constructed directional liftover chains using nf-LO (*62*) (minimap2, -cx asm20 -f 0.02). Query and target chromosomes were split into 10-Mb segments with 100-kb overlaps before alignment. Internal unchained gaps ≥500 bp were subjected to local rescue using 50-kb flanking sequences. Gaps > 200 kb were divided into ≤ 200-kb windows and subjected to sequential rescue alignments using minimap2 (asm20), MUMmer4 (--maxmatch), and LASTZ (-- identity=40 --coverage=1 --hspthresh=300 --gappedthresh=300 --ydrop=10000). Rescued alignments were merged with backbone chains and filtered using UCSC chainPreNet and chainNet (-minScore=10000) to generate the final directional chains.

### Evaluation of cross-genome coordinate conversion

We evaluated coordinate conversion using chromosomes 1B, 2A, 3D and 6A from ten wheat assemblies (AK58 (*38*), IWGSC RefSeq v1.0 (*4*), Jagger (*12*), Jin50 (*32*), JM22 (*11*), Kariega (*34*), KN9204 (*41*), LongReach Lancer (*12*), JM44 (*63*) and YM33 (*33*)). For each chromosome, ten independent sets of 1,000 non-redundant genomic sites were randomly sampled. Only sites with a complete 201-bp flanking sequence were retained. Sites mapping to exactly one target coordinate were considered uniquely converted. We calculated the conversion rate as the proportion of uniquely converted sites. For accuracy, we extracted the corresponding 201-bp target sequences, reverse-complementing minus-strand mappings, and calculated sequence identity via Levenshtein distance. Conversions with ≥95% identity were deemed accurate, and coordinate accuracy was calculated as the proportion of accurate conversions among all assessable uniquely converted sites.

### Identification and gene-order clustering of NLR loci

We identified NLR loci using NLR-Annotator v2.1b (*64*). Candidate loci were extracted with 20-kb flanks and subjected to gene prediction with AUGUSTUS (*65*) using the wheat model (--species=wheat, --strand=both and --genemodel=partial). Predicted proteins were screened for the NB-ARC domain (PF00931) using HMMER (*66*), retaining models with a domain conditional E-value ≤ 1 × 10^−5^. For sequence-level non-redundancy, PF00931-supported CDSs were clustered with CD-HIT-EST (*67*) at 95% nucleotide identity and ≥ 90% coverage of the shorter sequence.

For locus clustering, NLR loci were placed within the gene order of the gene annotation. CS-NP chromosomes were treated as continuous segments, while non-CS pangenomes were partitioned by original donor blocks. Loci without overlapping genes were inserted as pseudo-genes. Pairs within the same biological segment were connected if their gene-order rank difference was ≤ 5, with edge weights defined as the inverse of genomic midpoint distance. NLR communities were identified using the Walktrap algorithm (*68*) implemented in python-igraph (*69*) with four-step random walks, defining clusters as communities with at least two loci.

### Gene Ontology enrichment of non-CS genes

For each WheatPanRef gene, the longest protein isoform was annotated using eggNOG-mapper (*70*) to obtain Gene Ontology (GO) terms. Non-CS genes were used as the foreground, and all annotated WheatPanRef genes as the background. Direct GO annotations were propagated to ancestral terms using GO.db. Over-representation analysis was performed with clusterProfiler (*71*), testing gene sets containing 5-5,000 genes and adjusting P-values via the Benjamini-Hochberg procedure (FDR ≤ 0.05). For visualization, we retained significant Biological Process terms with fold enrichment > 1, ≥ 10 foreground genes, and GO depth 3-8, excluding overly broad terms (> 15% of foreground genes).

### Whole-genome resequencing and variant calling

We processed WGS reads from 485 bread wheat accessions (*72*) using a standard BWA-GATK pipeline **(table S6)**. Raw reads were trimmed with Trimmomatic (v0.38) (*73*) and aligned to WheatPanRef with BWA-MEM (*26*). The resulting alignments were sorted and indexed with Samtools (v1.31) (*43*). Duplicate reads were marked using the MarkDuplicates module of Picard (v2.24.0, https://broadinstitute.github.io/picard). Single-nucleotide polymorphisms (SNPs) and short insertions/deletions (InDels) were discovered with the HaplotypeCaller module of GATK (v4.1.8.0) (*31*) in GVCF mode, calling variants per sample across genomic intervals in parallel (using the -ERC GVCF and -L interval options). The per-sample GVCF files were subsequently consolidated and jointly genotyped with the GenotypeGVCFs module to produce a cohort-level raw variant call set. Raw calls were subjected to hard filtering following GATK’s best-practice recommendations: SNPs with a quality by depth (QD) < 2.0, Fisher strand (FS) > 60.0, strand odds ratio (SOR) > 3.0, mapping quality (MQ) < 40.0, MQRankSum < −12.5, and ReadPosRankSum < −8.0 were removed. InDels were filtered with the more stringent thresholds (QD < 2.0, FS > 200.0, SOR > 10.0, and ReadPosRankSum < −20.0). The filtered set was further refined by removing sites with high missingness and non-biallelic sites, and by excluding variants overlapping repetitive or low-complexity regions. Mapped-read counts were obtained for each accession from the BAM files using Samtools (v1.31) (*43*) to get the number of mapped reads. The SNP density was calculated in 1-Mb bins, and the nucleotide diversity was calculated in sliding windows of 10-kb per chromosome using VCFtools (v0.1.17) (*74*).

### CNV-based association analysis and local haplotype characterization

Copy-number variations (CNVs) were identified from 10-kb read counts obtained using SAMtools (*43*) and BEDTools (*45*) (properly paired reads, MAPQ ≥ 10) a (properly paired reads, MAPQ ≥ 10) across the 485 accessions (*72*). Read counts were normalized per kilobase to the median non-zero depth of the CS backbone, and adjacent windows with similar patterns (Jaccard similarity ≥ 0.9) were merged into CNV markers. To account for population structure, biallelic CS-NP SNPs were filtered for minor allele frequency ≥ 0.05 and missingness ≤ 5%, followed by linkage disequilibrium pruning (1-Mb window, r² ≤ 0.2) using PLINK 2.0 (*75*). Genome-wide CNV associations were tested using GEMMA (*76*) linear mixed models with ten SNP-derived principal components and population groups as covariates, alongside a leave-one-chromosome-out CNV kinship matrix. Associations with *P*-values < 1 × 10⁻⁶ were considered significant.

To identify alternative sequences associated with CS deletions, we evaluated normalized read-depth support for non-CS blocks in deletion carriers versus non-carriers. Local SNP association analysis was performed within candidate non-CS regions using GEMMA (*76*), retaining samples with ≥ 90% local genotype call rates and SNPs with missingness ≤ 10% and minor allele count ≥ 5. Local significance was determined via Bonferroni correction. Haplotypes were defined from peak-associated SNP genotypes, and thousand-grain weight differences were assessed using two-sided Wilcoxon rank-sum tests.

## Data availability

This unique resource is made available in open access by China Agricultural University in a dedicated website (https://wheat.cau.edu.cn/WheatPanRef) and to the entire wheat community through the WheatIS Data Discovery following the FAIR principles (https://urgi.versailles.inrae.fr/wheatis/).

