## Supplementary Figures and Legends for "WheatPanRef: a unified linear pan-reference genome for wheat genomics"

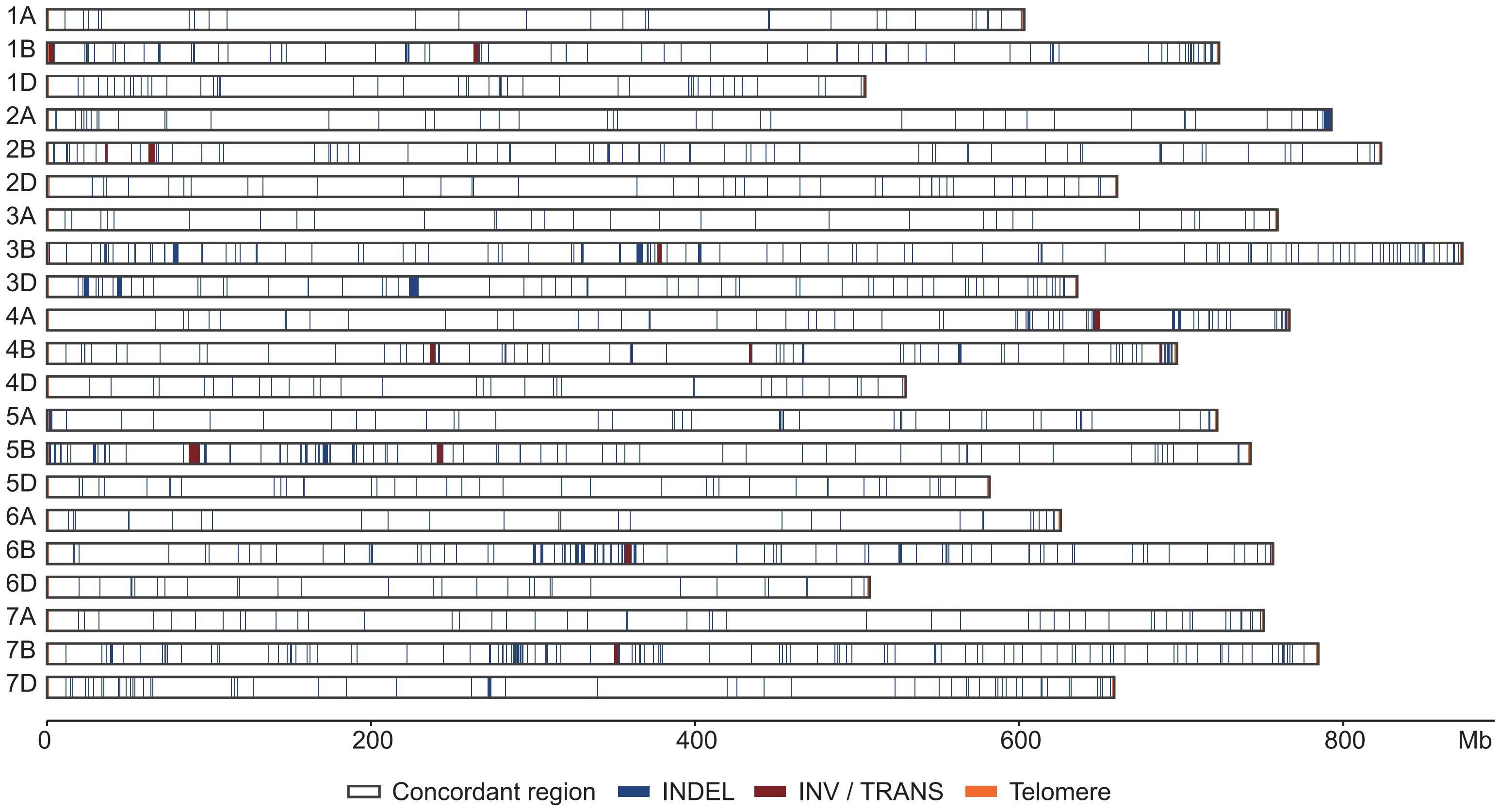


**fig S1 | Chromosome lengths and distribution of assembly-discrepant regions in CS-NP.**
White, blue, dark-red and orange regions represent the concordant segments, InDels, inversions or translocations, and telomeric sequences, respectively.

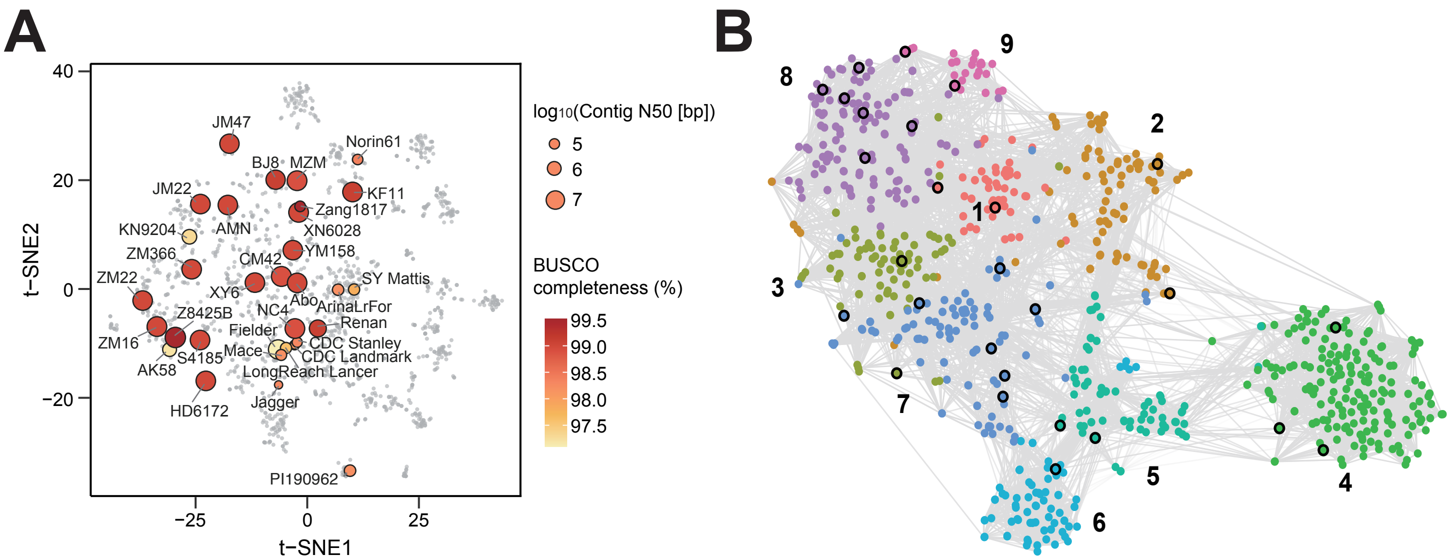


**fig S2 | Genetic diversity and assembly quality of WheatPanRef donor genomes.**
**(A)** t-distributed stochastic neighbour embedding (t-SNE) of wheat accessions based on genome-wide SNPs. The 37 donor accessions are labelled, with circle size indicating contig N50 and colour indicating BUSCO completeness. Other accessions are shown in grey.
**(B)** Germplasm resource-based identity-by-descent (gIBD) similarity network showing nine genetic groups. Nodes represent accessions, colours indicate genetic groups and edges denote pairwise genetic similarity. Black outlines highlight the 37 donor accessions.

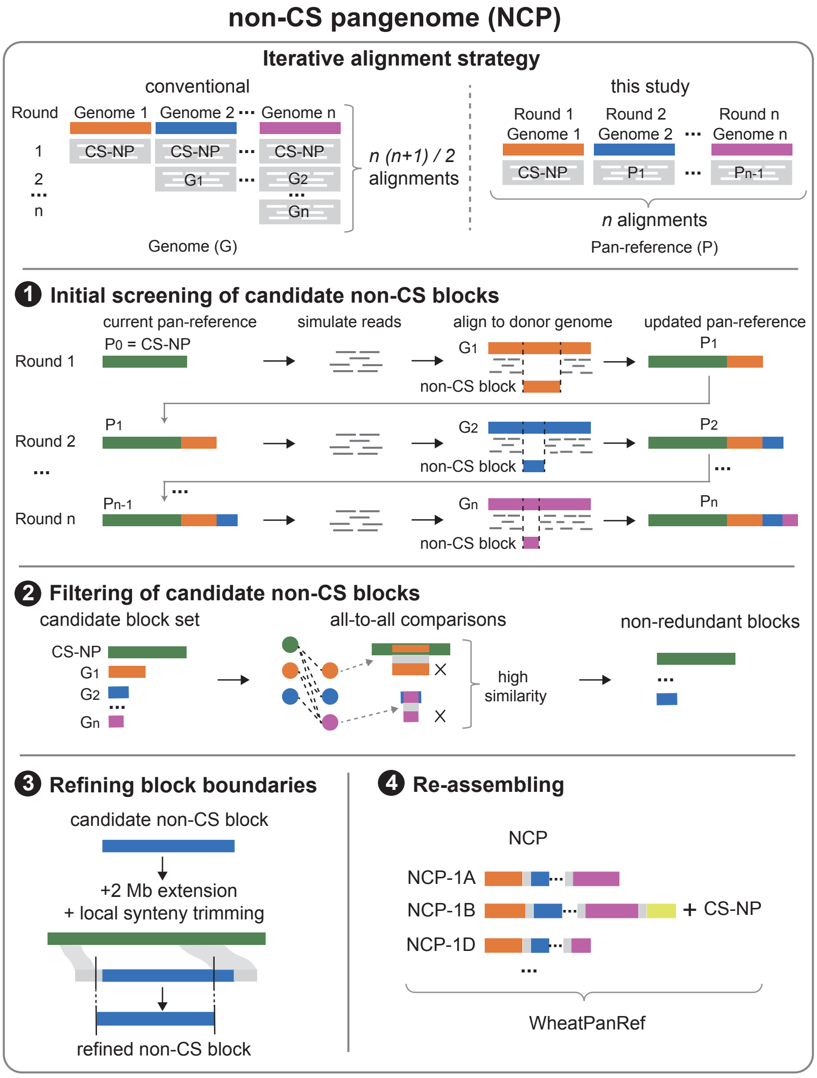


**fig S3 | Iterative construction of the non-CS pangenome.**
Conventional genome comparisons require $n(n + 1)/2$ alignments, whereas the iterative map-to-pan strategy requires only $n$ primary alignments. Chinese Spring reads were initially mapped to the first donor assembly, followed by simulated reads from the expanding pan-reference mapped to successive donors to identify non-CS blocks. Candidate blocks were filtered for redundancy, refinedby local synteny and concatenated by chromosome with 500-bp N spacers to generate the non-CS pangenome.

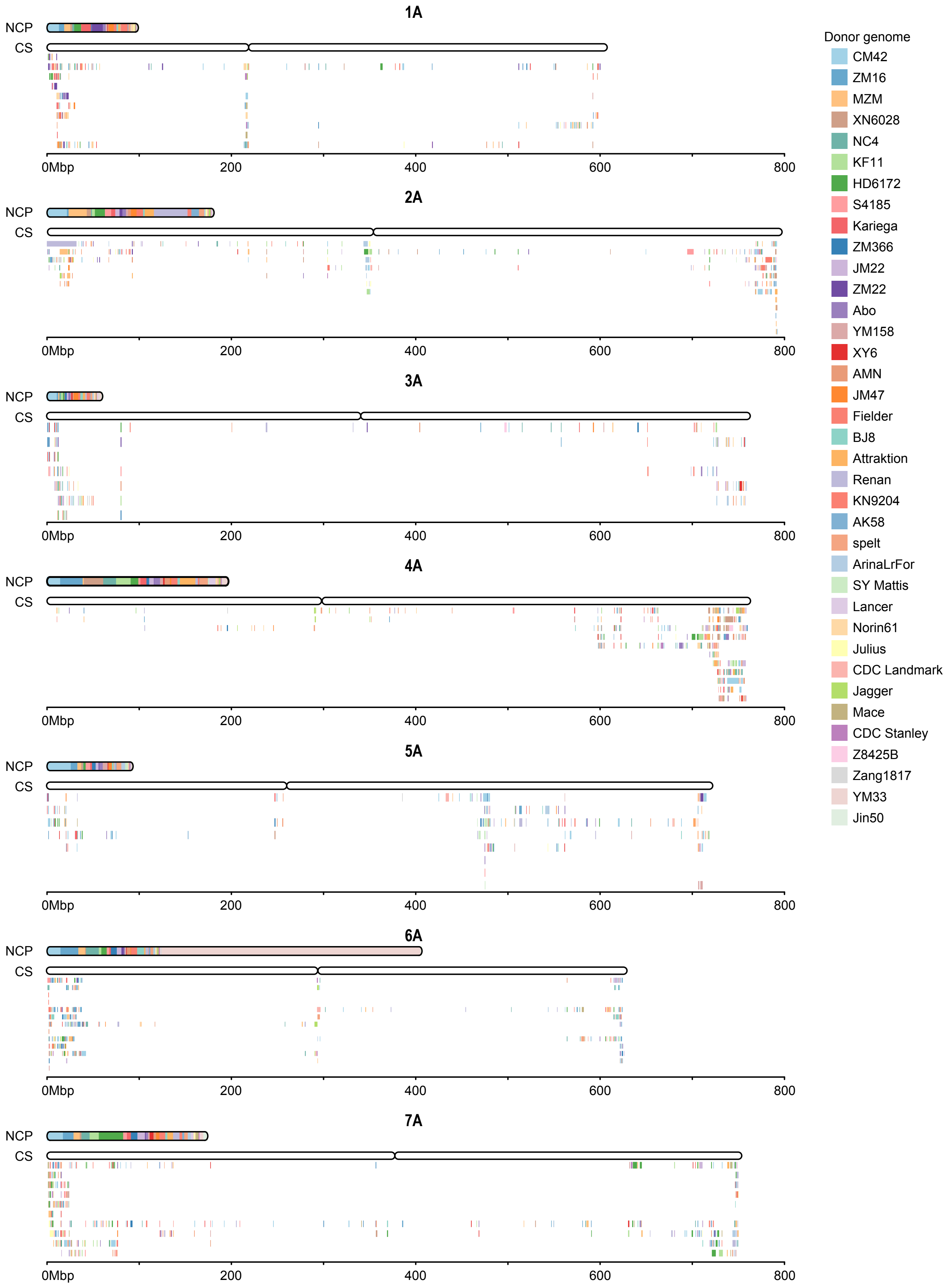


**fig S4 | Composition and chromosomal distribution of non-CS sequences in the A subgenome.**
The upper bars show the donor composition of non-CS pseudochromosomes (NCPs), and the lower tracks show their homologous intervals on CS-NP chromosomes. Overlapping intervals are stacked vertically. Colours indicate donor genomes.

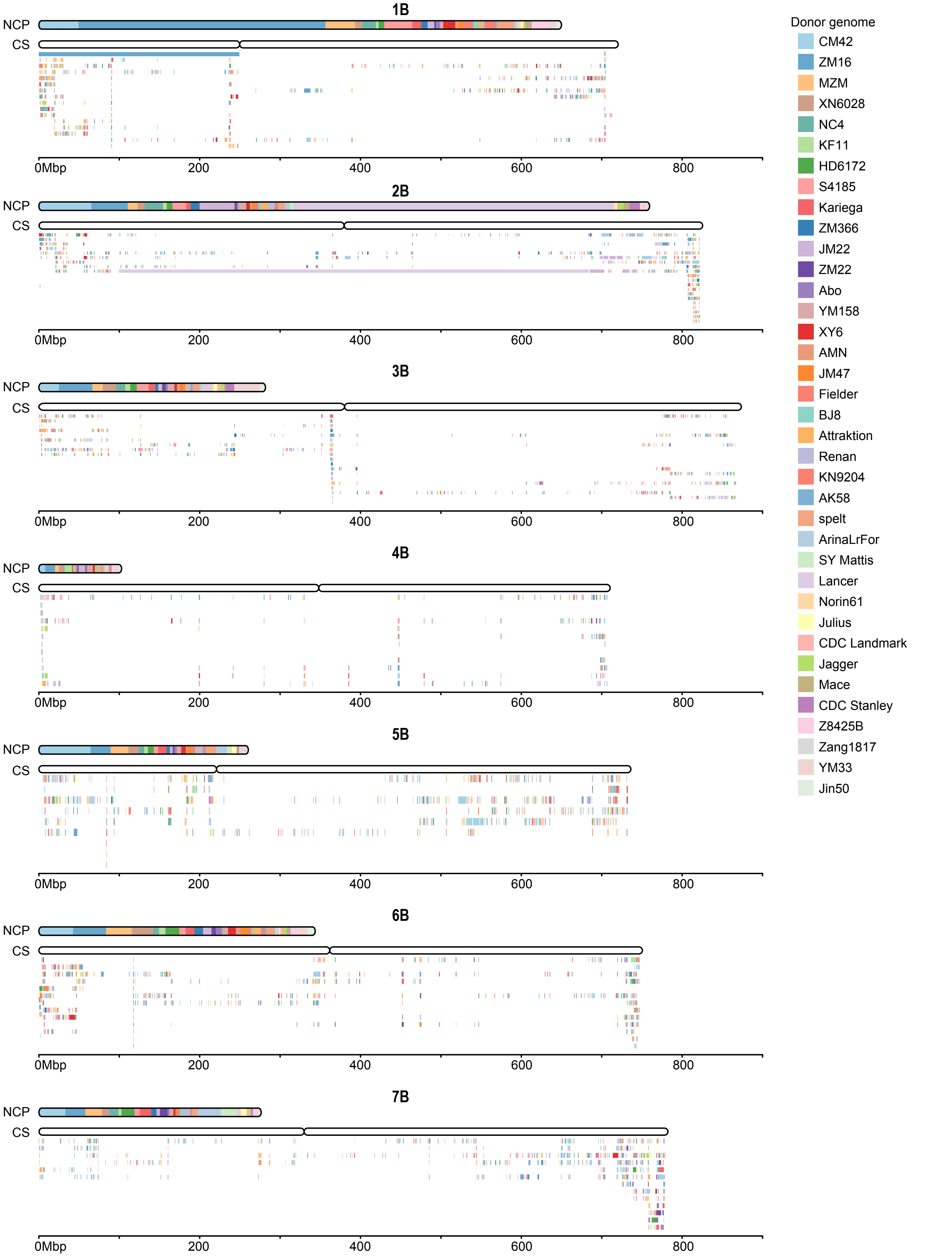


**fig S5 | Composition and chromosomal distribution of non-CS sequences in the B subgenome.**
The upper bars show the donor composition of non-CS pseudochromosomes (NCPs), and the lower tracks show their homologous intervals on CS-NP chromosomes. Overlapping intervals are stacked vertically. Colours indicate donor genomes.

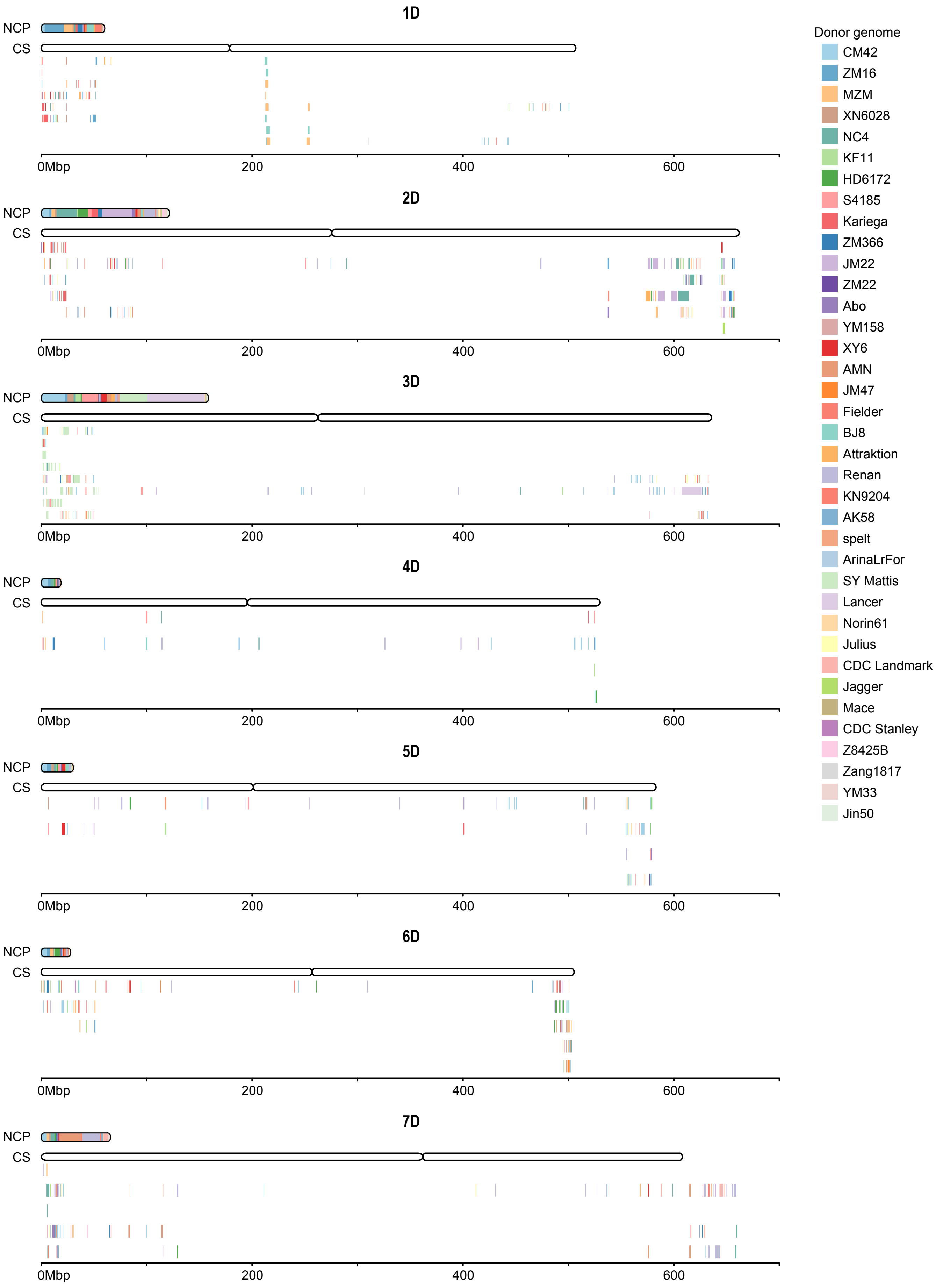


**fig S6 | Composition and chromosomal distribution of non-CS sequences in the D subgenome.**
The upper bars show the donor composition of non-CS pseudochromosomes (NCPs), and the lower tracks show their homologous intervals on CS-NP chromosomes. Overlapping intervals are stacked vertically. Colours indicate donor genomes.

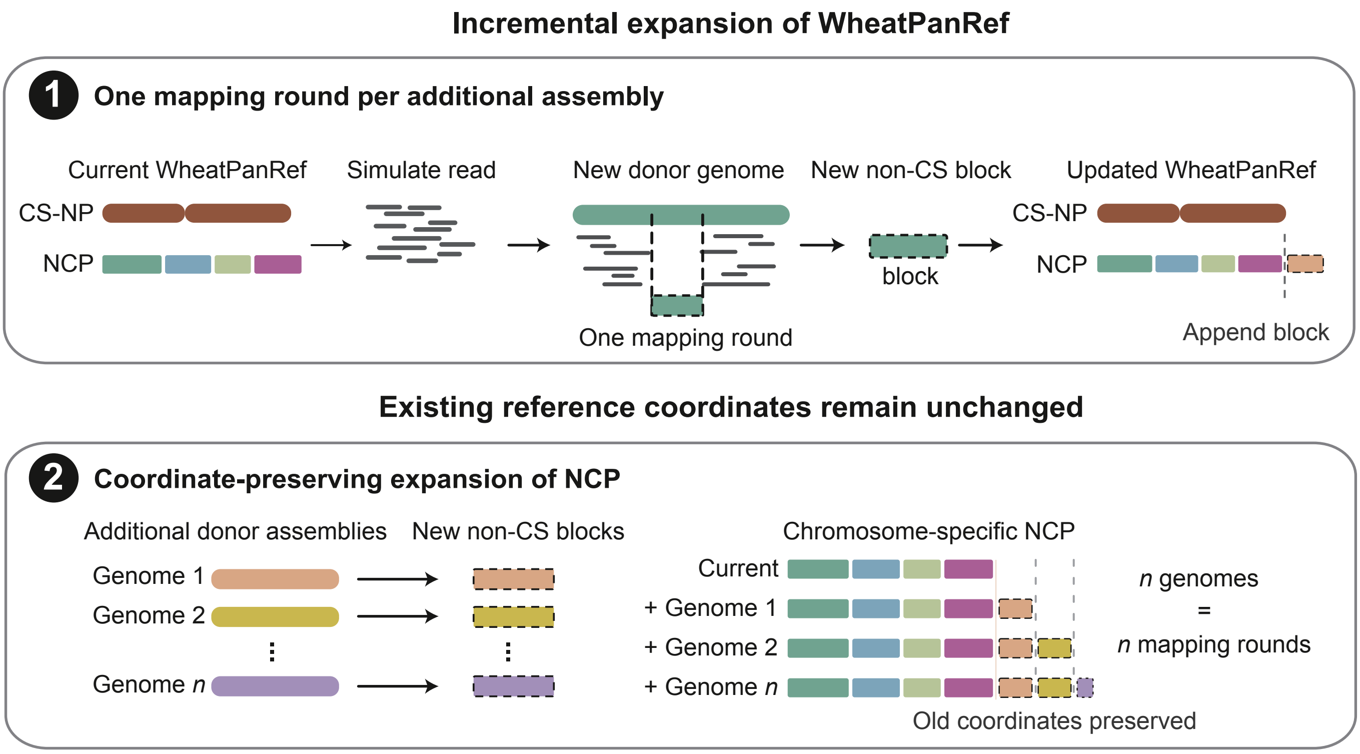


**fig S7 | Incremental expansion of WheatPanRef.**
Simulated reads from the current WheatPanRef are mapped to each new donor assembly to identify novel non-CS blocks, which are appended to the corresponding NCP chromosomes without altering existing coordinates. Incorporating *n* additional assemblies thus requires only *n* mapping rounds.

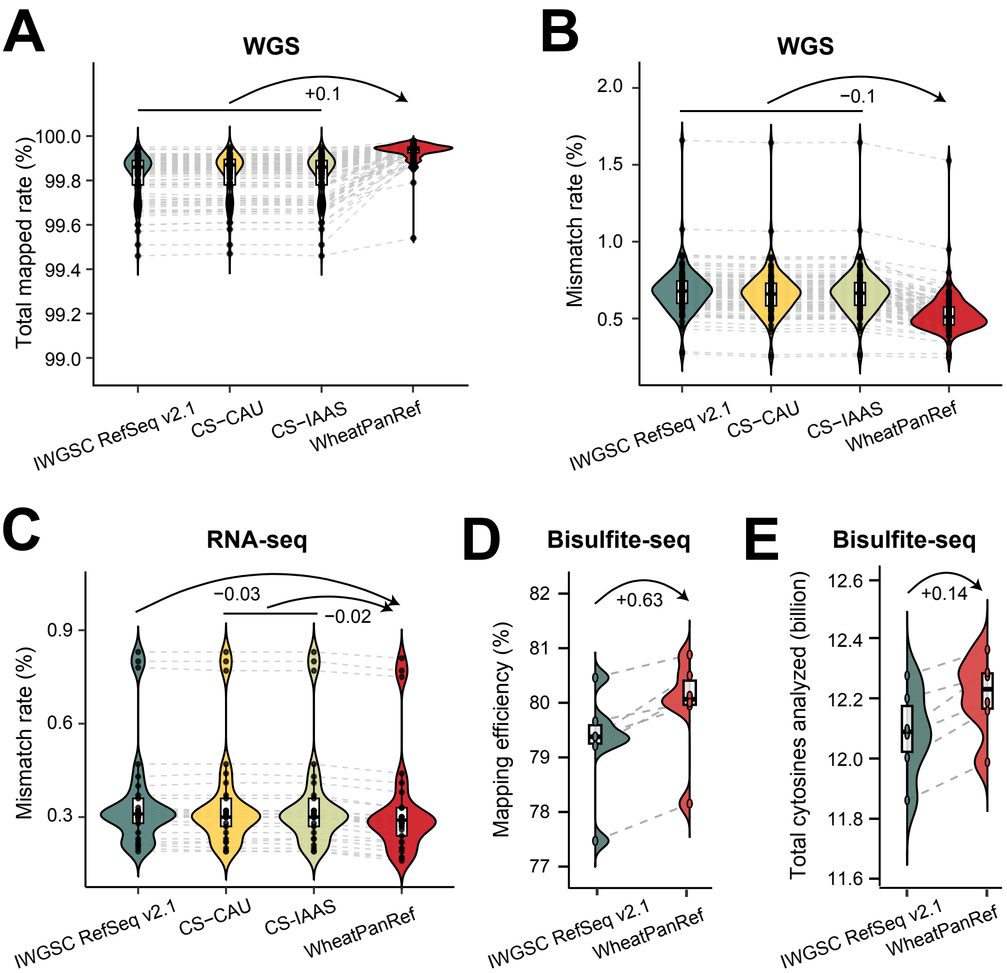


**fig S8 | Improved read-alignment performance of WheatPanRef across multiple sequencing datasets.**

**(A,B)** Total mapping rates (A) and mismatch rates (B) for whole-genome resequencing reads from 100 wheat accessions aligned to IWGSC RefSeq v2.1, CS-CAU, CS-IAAS and WheatPanRef.

**(C)** Mismatch rates for 29 RNA-seq datasets aligned to the same four references.

**(D, E)** Mapping efficiency (D) and total cytosines analysed (E) for six bisulfite-sequencing datasets aligned to IWGSC RefSeq v2.1 and WheatPanRef. Grey dashed lines connect measurements from the same sample; arrows indicate differences in mean values.

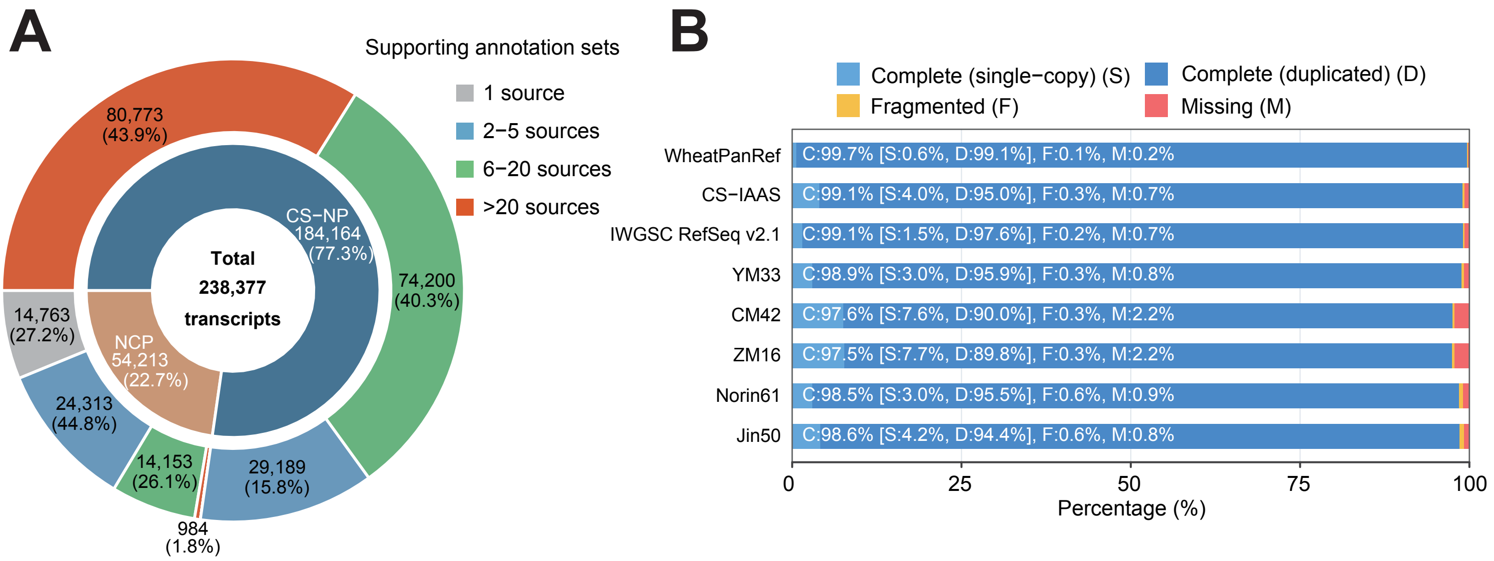


**fig S9 | Cross-source support and completeness of WheatPanRef gene annotations.**
**(A)** Genomic distribution and cross-annotation support of 238,377 WheatPanRef transcripts. The inner ring shows the proportions of transcripts on CS-NP and the non-CS pangenome (NCP), and the outer ring shows their distribution by the number of supporting annotation sets (1, 2–5, 6–20 and >20). Two single-source CS-NP transcripts are not separately labeled.
**(B)** BUSCO assessment of protein-coding gene annotation completeness in WheatPanRef and seven wheat genome assemblies using the poales_odb12 dataset (n = 6,282). Bars indicate complete single-copy (S), complete duplicated (D), fragmented (F) and missing (M) BUSCOs; C denotes total completeness.

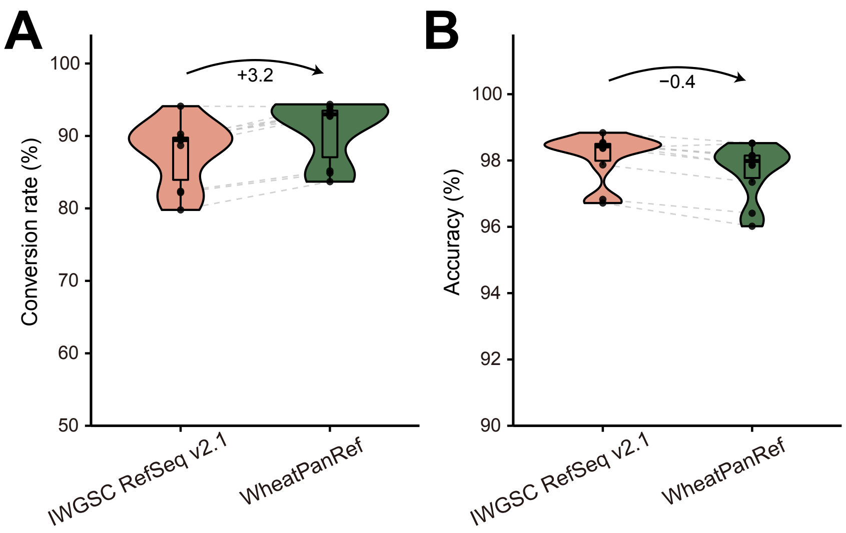


**fig S10 | Evaluation of WheatPanRef-mediated cross-genome coordinate conversion.**
**(A, B)** Conversion rate (A) and accuracy (B) for coordinates transferred from ten wheat assemblies to Fielder via IWGSC RefSeq v2.1 or WheatPanRef. Black dots represent assembly-level means, grey dashed lines connect measurements from the same assembly, and arrows indicate mean differences in percentage points.

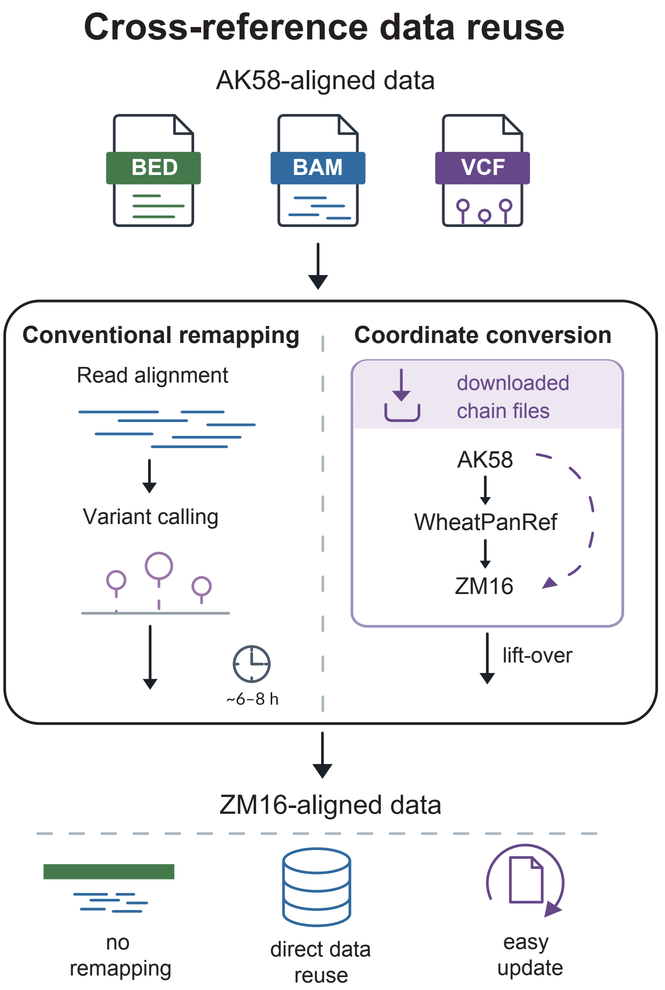


**fig S11 | Offline coordinate transfer and genomic data reuse through WheatPanRef.**
Schematic comparison of conventional remapping and chain file-mediated transfer of BED, BAM and VCF data from AK58 to ZM16 through WheatPanRef, enabling cross-reference data reuse without read remapping.

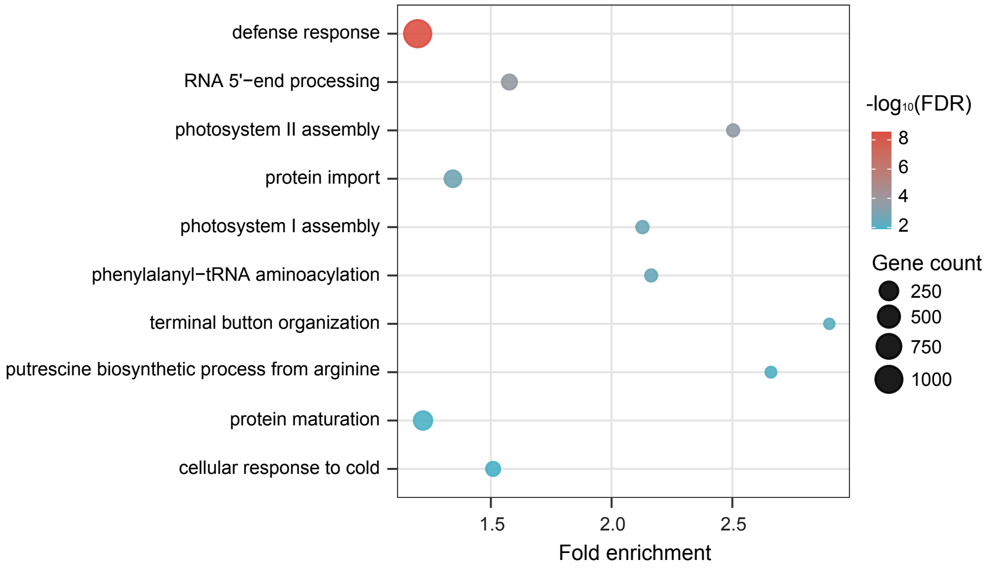


**fig S12 | Gene Ontology enrichment of non-CS pangenome genes.**
Representative enriched Gene Ontology (GO) biological-process terms among non-CS genes, using all GO-annotated WheatPanRef genes as the background. Bubble size indicates gene count, and colour represents enrichment significance ($-\log_{10}$FDR; Benjamini-Hochberg adjusted *P* value).

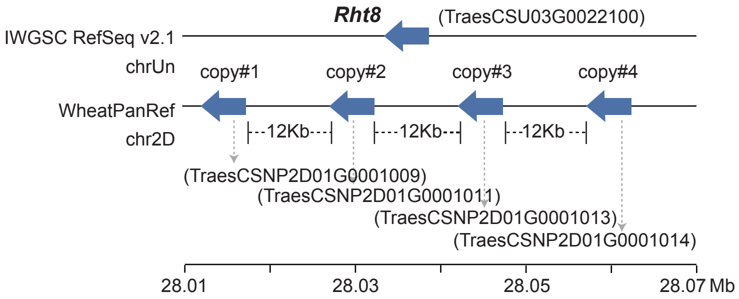


**fig S13 | Chromosomal anchoring and copy-number resolution of *Rht8* in WheatPanRef.**
The *Rht8* locus (TraesCSU03G0022100), unplaced in IWGSC RefSeq v2.1, was anchored to CS-NP chromosome 2D (28.01-28.07 Mb), revealing a four-copy tandem array with approximately 12-kb intergenic spacers.

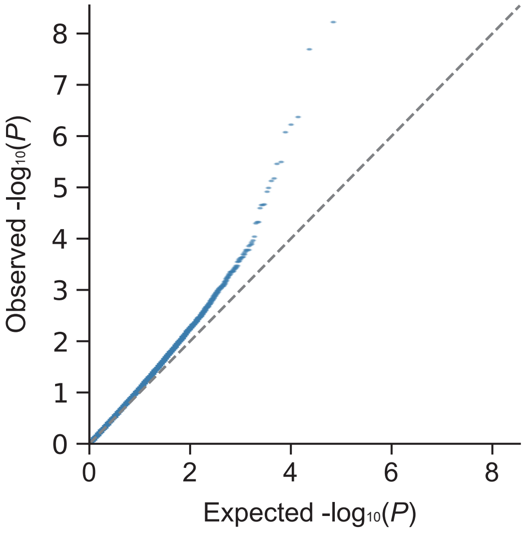


**fig S14 | Quantile–quantile plot of CNV associations with thousand-grain weight.**
Observed versus expected $-\log_{10}(P)$ values from genome-wide association analysis of 34,198 copy-number variation (CNV) markers in 485 wheat accessions. The dashed line indicates the null expectation.
